# A connectomics-based taxonomy of mammals

**DOI:** 10.1101/2022.03.11.483995

**Authors:** Laura E. Suárez, Yossi Yovel, Martijn P. van den Heuvel, Olaf Sporns, Yaniv Assaf, Guillaume Lajoie, Bratislav Misic

## Abstract

Mammalian taxonomies are conventionally defined by morphological traits and genetics. How species differ in terms of neural circuits and whether inter-species differences in neural circuit organization conform to these taxonomies is unknown. The main obstacle for the comparison of neural architectures have been differences in network reconstruction techniques, yielding species-specific connectomes that are not directly comparable to one another. Here we comprehensively chart connectome organization across the mammalian phylogenetic spectrum using a common reconstruction protocol. We analyze the mammalian MRI (MaMI) data set, a database that encompasses high-resolution *ex vivo* structural and diffusion magnetic resonance imaging (MRI) scans of 124 species across 12 taxonomic orders and 5 superorders, collected using a single protocol on a single scanner. We assess similarity between species connectomes using two methods: similarity of Laplacian eigenspectra and similarity of multiscale topological features. We find greater inter-species similarities among species within the same taxonomic order, suggesting the connectome organization recapitulates traditional taxonomies defined by morphology and genetics. While all connectomes retain hallmark global features and relative proportions of connection classes, inter-species variation is driven by local regional connectivity profiles. By encoding connectomes into a common frame of reference, these findings establish a foundation for investigating how neural circuits change over phylogeny, forging a link from genes to circuits to behaviour.

## INTRODUCTION

Anatomical projections between brain regions form a complex network of polyfunctional neural circuits [86]. Signaling on the brain’s connectome is thought to support cognition and the emergence of adaptive behaviour. Advances in imaging technologies have made it increasingly feasible to reconstruct the wiring diagram of biological neural networks. Thanks to extensive international data sharing efforts, these detailed reconstructions of the nervous system’s connection patterns have been made available in humans and in multiple model organisms [95], including invertebrate [25, 94, 102, 104], avian [83], rodent [20, 71, 78], feline [18, 33, 79] and primate species [52, 55, 57].

The rising availability of connectomics data facilitates cross-species comparative studies that identify commonalities in brain network topology and universal principles of connectome evolution [10, 11, 95]. A common thread throughout these studies is the existence of non-random topological attributes that theoretically enhance the capacity for information processing [85]. These include a highly clustered architecture with segregated modules that promote specialized information processing [43, 101], as well as a densely interconnected core of high-degree hubs that shortens communication pathways [96], promoting the integration of information from distributed specialized domains [5, 108]. These universal organizational features suggest that connectome evolution has been shaped by two opposing and competitive pressures: maintaining efficient communication while minimizing neural resources used for connectivity [23].

While comparative analysis can focus on commonalities among mammalian connectomes and identify universal wiring principles, it can also be used to systematically explore differences among connectomes that confer specific adaptive advantages. Indeed, despite commonalities, architectural variations are also observed even among closely related species [10]. Factors such as the external environment, genetics and distinct gene expression programs also account for diversity in neural connectivity patterns [63]. Subtle variations in connectome organization may potentially account for species-specific adaptations in behaviour and cognitive function.

But how does the connectome vary over phylogeny? Traditionally, mammalian taxonomies were built on morphological differences among species [29]. Besides physical commonalities, species within the same taxonomic group also tend to share similar behavioural repertoires [17, 105, 106]. Modern high-throughput whole-genome sequencing has further delineated phylogenetic links and relationships among mammalian species [1, 28, 67, 81]. In addition to refining the overall classification of mammals, whole-genome comparative analyses have established the genetic basis of phenotypic variation across phylogeny [67]. Whether inter-species differences in the organization of connectome wiring conform to this taxonomy remains unknown. How do genes sculpt behaviour across evolution? Could speciation events in the genome leading to variations in connectome architecture be the missing link between genomics and behaviour? Rigorously addressing these questions is challenging due to the lack of methodological consistency in the acquisition and reconstruction of neural circuits, or the limited number of available species.

Here we comprehensively chart connectome organization across the mammalian phylogenetic spectrum. We analyze the mammalian MRI (MaMI) data set, a comprehensive database that encompasses high-resolution *ex vivo* diffusion and structural (T1- and T2-weighted) MRI scans of 124 species (a total of 225 scans including replicas) [3]. All images were acquired using the same scanner and protocol. All connectomes were reconstructed using a uniform parcellation scheme consisting of 200 brain areas, including cortical and subcortical regions. Notably, the MaMI data set spans a wide range of categories across different taxa levels of morphologic and phylogenetic mammalian taxonomies [3]. Specifically, it includes animal species across 5 different superorders (Afrotheria, Euarchontoglires, Laurasiatheria, Xenarthra and Marsupialia) and 12 different orders (Cetartiodactyla, Carnivora, Chiroptera, Eulipotyphla, Hyracoidea, Lagomorpha, Marsupialia, Perissodactyla, Primates, Rodentia, Scandentia and Xenarthra).

Taking advantage of the harmonized imaging and reconstruction protocols, we quantitatively assess the similarity of species’ connectomes to construct data-driven phylogenetic relationships based on brain wiring. We compare these inter-species wiring similarities with conventional morphologically and genetically defined mammalian taxonomies. We determine the extent to which connectome topology conforms to established taxonomic classes, and identify network features that are associated with speciation.

## RESULTS

The MaMI data set consists of high-resolution *ex vivo* diffusion and structural (T1- and T2-weighted) MRI scans of 124 species. Since there is no species-specific template, all connectomes were reconstructed using a uniformly applied 200-node parcellation. Having equally sized networks facilitates graph comparison but also implies a lack of direct correspondence between nodes across species. However, because our focus is on the statistics of connectomes’ topology, this does not impact our analyses. As the size of the network is kept constant across all species, voxel size is normalized to brain volume. Fig. 1 shows the distribution of connectomes across ten mammalian orders (out of the twelve present in the data set). We focus on the six orders that contain 5 or more distinct species (within the Laurasiatheria and Euarchontoglires superorders); these include Chiroptera, Rodentia, Artiodactyla, Carnivora, Perissodactyla and Primates, resulting in a total of 111 different animal species and 203 brain scans. A complete list of the animal species included in the analyses is provided in Table S1.

**FIG. 1.**
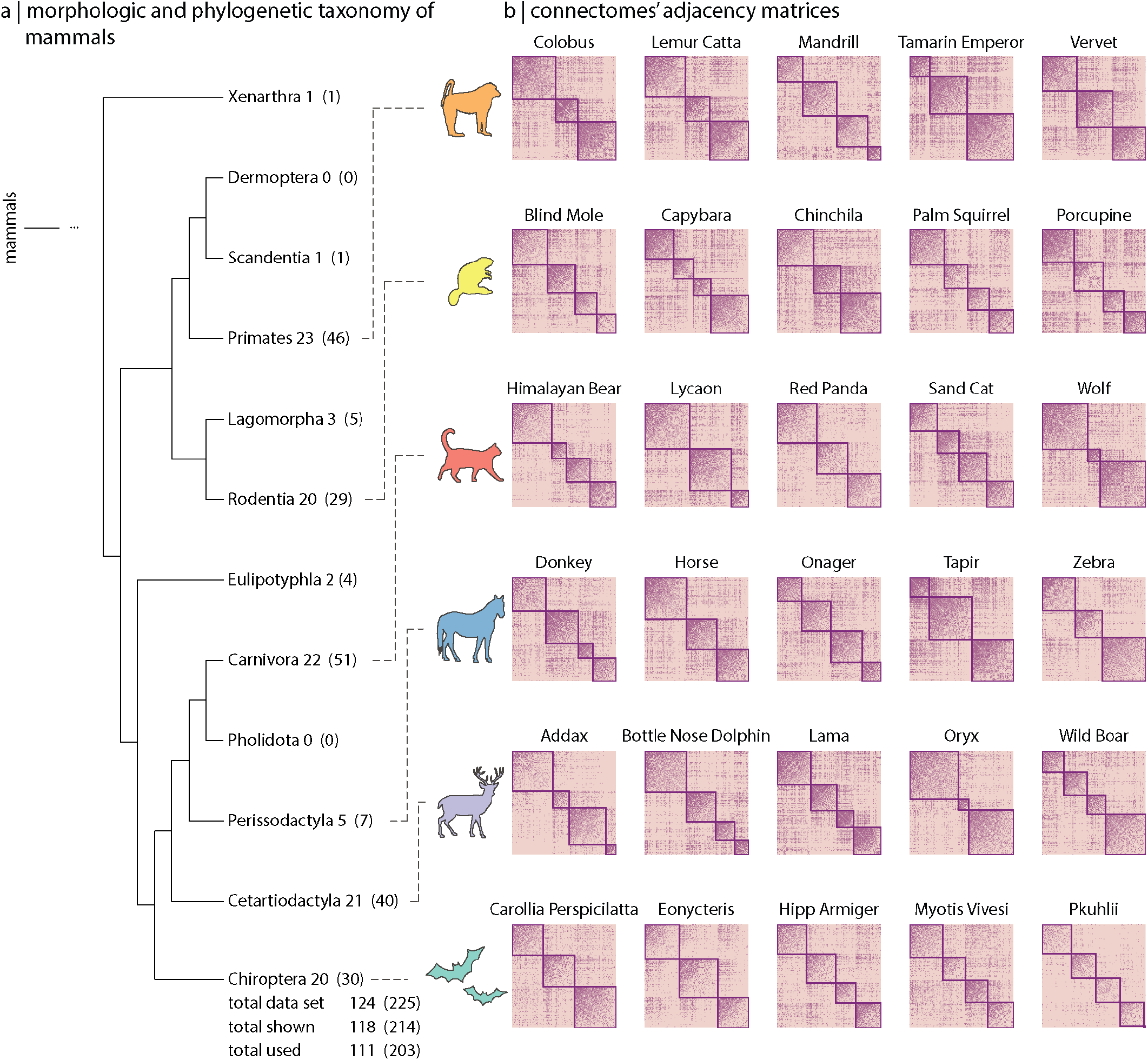
MaMI data set. The MaMI data set encompasses high-resolution *ex vivo* structural and diffusion MRI scans of 124 animal species spanning 12 morphologically and phylogenetically defined taxonomic orders: Cetartiodactyla, Carnivora, Chiroptera, Eulipotyphla, Hyracoidea, Lagomorpha, Marsupialia, Perissodactyla, Primates, Rodentia, Scandentia and Xenarthra. (a) Hierarchical relationships across 10 (out of the 12 included in the data set) morphologic and phylogenetic taxonomic orders. Numbers outside the parenthesis correspond to the number of unique species within each order and numbers inside the parenthesis correspond to the number of samples (including replicas). (b) Connectivity matrices for 5 randomly-chosen sample species within each of the six orders included in the analyses (i.e., Cetartiodactyla, Carnivora, Chiroptera, Perissodactyla, Primates and Rodentia). Only orders with at least 5 different species were included for the analyses. Nodes are organized according to their community affiliation obtained from consensus clustering applied on the connectivity matrix (see *Methods*). Communities in (b) correspond to the partition for which the resolution parameter *γ* = 1.0 (see Fig. S1).

### Connectome-based inter-species distances

Similarity between species’ network architectures is estimated using two network-based distance metrics: spectral distance, based on the eigenspectrum of the normalized Laplacian of the connectivity matrix (see Fig. S2; [31]), and topological distance, based on a combination of multiscale graph features of the binary and weighted connectivity matrices (Fig. S3 and Fig. S4 show the distribution of individual local and global graph features, respectively; [77]). For completeness, Fig. S5 and Fig. S6 show the cumulative distribution of binary and weighted local features, respectively, for individual species. Both methods measure how similar the architectures of two connectomes are. By mapping connectomes to a common space, the normalized Laplacian eigenspectrum and the graph features of the connectivity matrix allow us to compare connectomes despite the fact that they come from different species, and that the nodes do not correspond to one another [59]. To account for the fact that some of the species have more than one scan, we randomly select one sample per specie and estimate (spectral and topological) inter-species distances. We repeat this procedure 10,000 times and report the average across iterations.

Fig. 2a shows the spectral distances between species’ connectomes. In general we observe smaller distances among members of the same order (outlined in yellow). Fig. 2b confirms this intuition by showing that spectral distances within orders (i.e., values along the diagonal) tend to be smaller than distances between orders (i.e., values off the diagonal). Fig. 2c shows the distributions of intra-and inter-order distances. The mean/median intra-order distance is significantly smaller than the mean/median inter-order distance (two-sample Wilcoxon-Mann-Whitney rank-sum test: median intra- and inter-order distances are 0.44 and 0.55, respectively, *P* < 10^−4^ two-tailed, and common-language effect size = 68%; two-sample Welch’s t-test: mean intra- and inter-order distances are 0.43 and 0.55, respectively, *P* < 10^−4^ two-tailed, and Cohen’s *d* effect size = 0.67; Fig. 2c). We find comparable results when estimating species similarity using topological distance (two-sample Wilcoxon-Mann-Whitney ranksum test: median intra- and inter-order distances are 0.41 and 0.53, respectively, *P* < 10^−4^ two-tailed, and common-language effect size = 66%; two-sample Welch’s t-test: mean intra- and inter-order distances are 0.41 and 0.53, respectively, *P* < 10^−4^ two-tailed, and Cohen’s *d* effect size = 0.59; Fig. 2d-f). These results suggest that species with similar genetics, morphology and behavior tend to have similar connectome architecture. In other words, variations in connectome architecture reflect phylogeny.

**FIG. 2.**
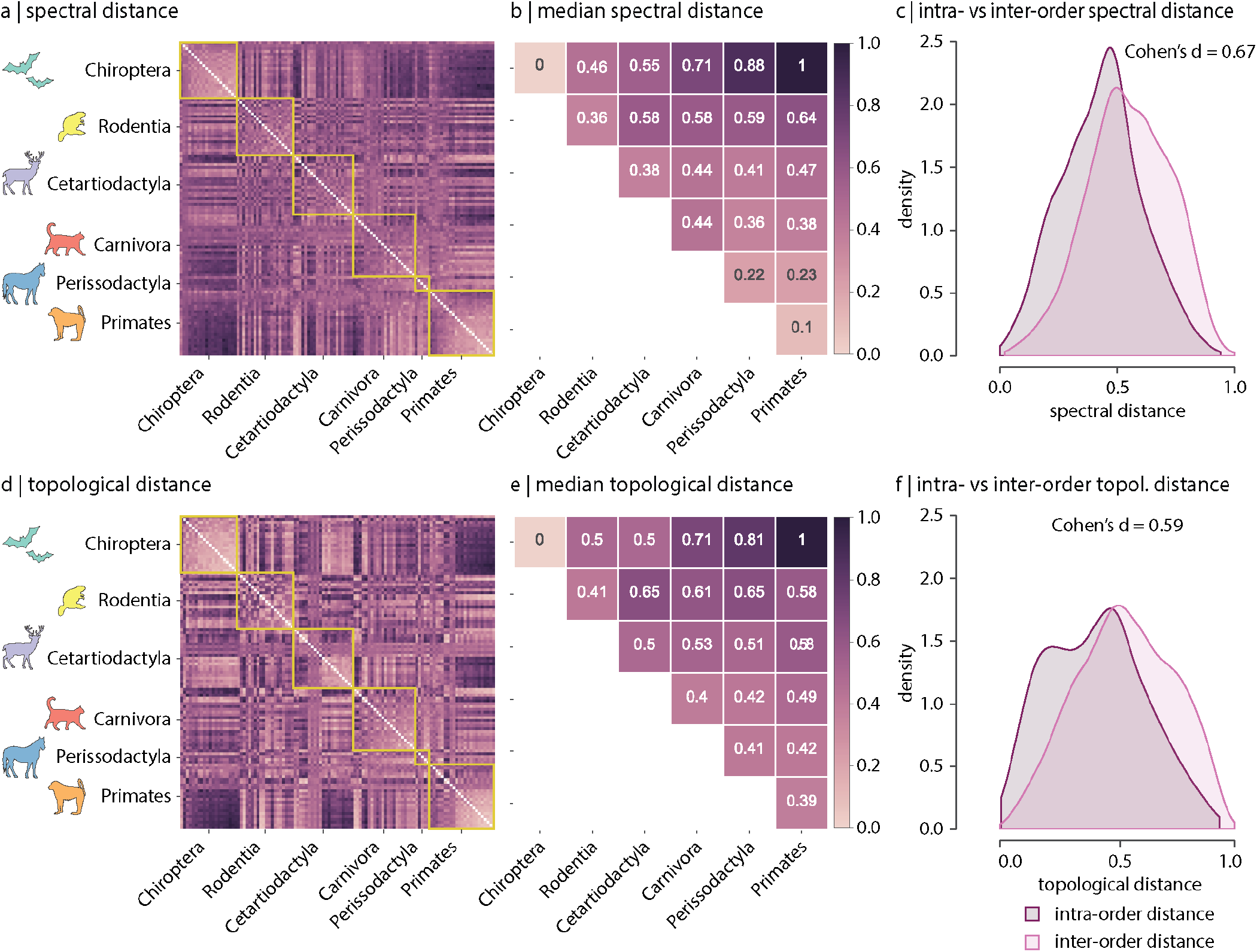
Spectral and topological distance between orders. (a) Spectral distance between species-specific connectomes. Lower distances indicate greater similarity. Yellow outlines indicate morphologically and genetically-defined orders. (b) Median spectral distance within and between all constituent members of each order. (c) Distribution of intra- and inter-order spectral distances. (d) Topological distance between species-specific connectomes. Lower distances indicate greater similarity. Yellow outlines indicate morphologically and genetically-defined orders. (e) Median topological distance within and between all constituent members of each order. (f) Distribution of intra- and inter-order topological distances. Effect sizes in (c) and (f) are the Cohen’s *d* estimator corresponding to a two-sample Welch’s t-test (*P* < 10^−4^). Equivalent conclusions are drawn if common-language effects sizes from the two-sample Wilcoxon-Mann-Whitney rank-sum test are used.

### Architectural features differentiate species

Next we consider which network features contribute to the differentiation (Fig. S3 and Fig. S4 show the distributions of local and global graph features, respectively). To address this question we recompute inter-species topo logical distances using different sets of graph features (Fig. 3). We find that the difference between intra- and inter-order topological distances tends to be larger when only local (node-level) features are included in the estimation of the topological distance (i.e., degree, clustering coefficient, betweenness and closeness; Fig. 3b,e), compared to when only global features are considered (i.e., characteristic path length, transitivity and assortativity; Fig. 3c,f). This is the case for both the binary and weighted versions of these features (top and bottom rows in Fig. 3, respectively). This suggests that differentiation of orders is better explained by differences in local network topology; conversely, global network topology appears to be conserved across species. A similar conclusion can be drawn when the eigenvalue distributions of the (normalized) Laplacian of the connectivity matrices are compared across species (Fig. S2). In spectral graph theory, the presence of eigenvalues with high multiplicities (e.g., *λ_i_* = 1) or eigenvalues equidistant to 1 provide information about the local organization resulting from recursive motif manipulation, such as the duplication of nodes or edge motifs (i.e., pairs of nodes) with the same connectivity profile [7, 8, 31]. The Laplacian eigenspectra of animal species belonging to different taxonomic orders tend to differ mostly around the eigenvalue 1, suggesting that differences across taxonomic orders are most likely due to the presence of different local connectivity fingerprints in the connectivity matrix (Fig. S2) [60, 61]. Consistent with this notion, the correlation between spectral and topological distance is maximum when only local binary features are included in the estimation of topological distance (Fig. S7).

**FIG. 3.**
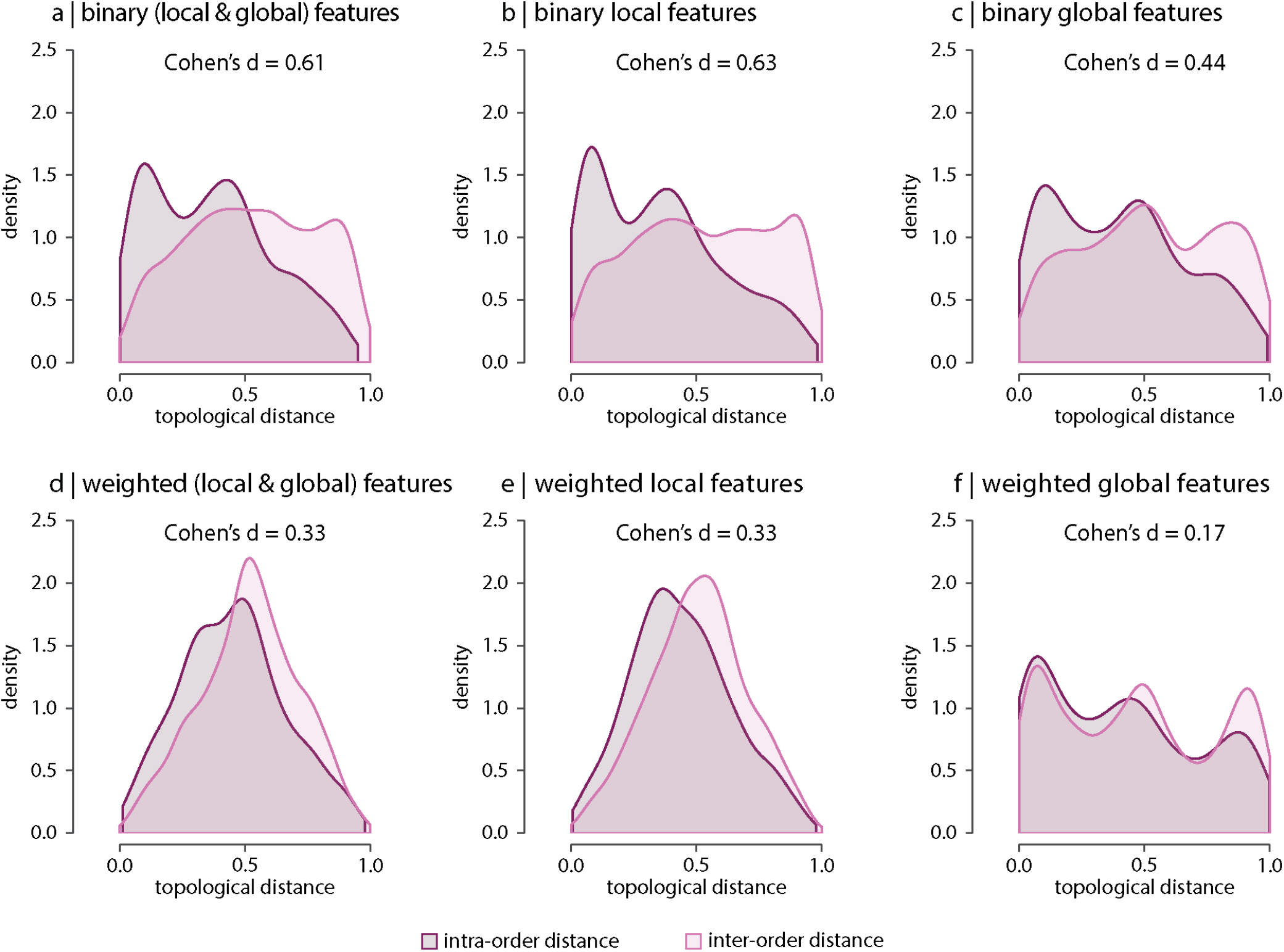
Contribution of network features. Topological distance can be computed using multiple local and global connectome features. Plots show intra- and inter-order distances when using only (a) binary local and global features, (b) binary local features, (c) binary global features, (d) weighted local and global features, (e) weighted local features and (f) weighted global features. Local features include (the average and standard deviation of) degree, clustering, betweenness, and closeness. Global features include characteristic path length, transitivity, and assortativity. For definitions please see *Methods*. Effect sizes correspond to the Cohen’s *d* estimator from a two-sample Welch’s t-test. Equivalent conclusions are drawn if common-language effects sizes from a two-sample Wilcoxon-Mann-Whitney rank-sum test are used. In all cases, the difference in the means and medians of intra- and inter-order distance distributions is statistically significant (*P* < 10^−4^). The same conclusions can be drawn after controlling for network density (Fig. S10).

We also observe that that the difference between intra- and inter-order topological distances is greater for binary than for weighted features (Fig. 3 a–c and d–f, respectively), independently of being local or global. This suggests that the strength of the connections is less important than the binary architecture of the connectivity matrix.

Some of the features used for the estimation of the topological distance depend on network density, which varies across taxonomic orders (Fig. S8). To determine whether the observed differences between intra- and inter-order distances are above and beyond differences due to network density, we perform the same analysis shown in Fig. 3, after controlling for density (Fig. S9). Results, shown in Fig. S10, suggest that differences between intra- and inter-order topological distances are not driven by differences in network density, but variations in wiring patterns, as captured by topological features, play a role in the observed phylogenetic variations in connec-tome organization.

### Conservation of small-world architecture

Anatomical brain networks are thought to simultaneously reconcile the opposing demands of functional integration and segregation by combining the presence of functionally specialized clusters with short polysynaptic communication pathways [12, 85, 86, 92]. Such architecture is often referred to as small-world, and is observed in a wide variety of naturally occurring and engineered networks [101]. Here we explore whether these principles of segregation and integration in global connectome organization are consistent across phylogeny. To do so, we estimate for each species the ratio of clustering coefficient to characteristic path length, normalized relative to a set of randomly rewired graphs that preserve the degree sequence of the nodes [44, 64, 77] (Fig. 4). Consistent with previous reports in individual species’ connectomes [12, 43, 87], we find that all connectomes display high and diverse levels of small-worldness, suggesting that simultaneously highly segregated and integrated networks is a global trait conserved across mammalian brains.

**FIG. 4.**
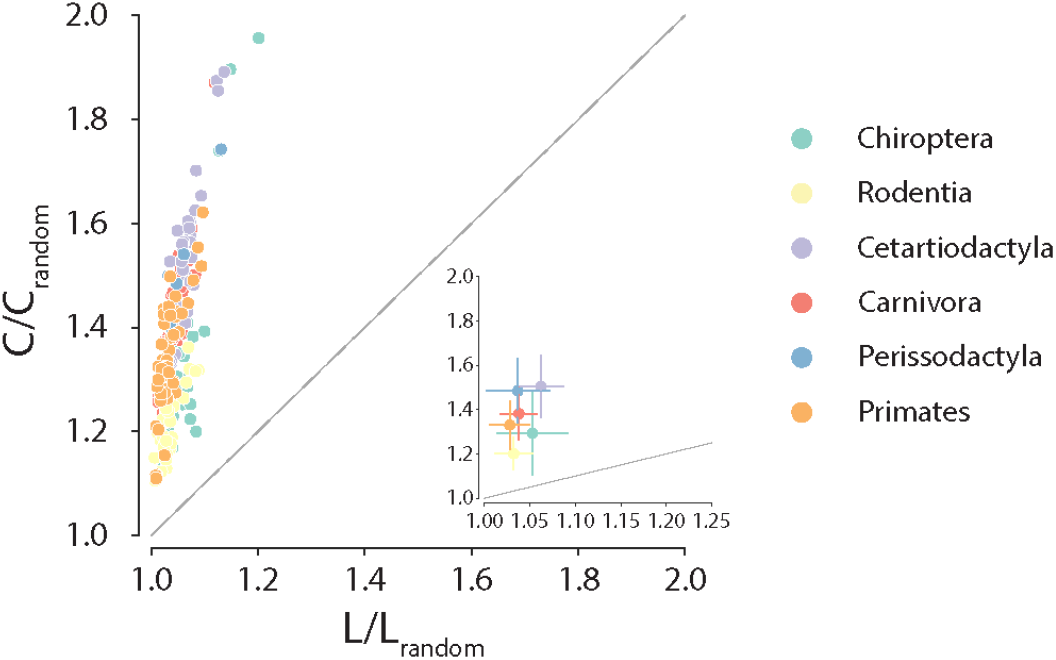
Conservation of small-world architecture. Clustering coefficient vs characteristic path length normalized relative to a set of 1000 randomly rewired graphs that preserve the degree sequence of the nodes [64]. For definitions of each graph measure, see *Methods*. Each data point represents a different animal species. Data points above the identity line are said to have small-world architecture. The inset on the right bottom corner is a zoom on the abscissa.

### Conservation of edge classes across species

The topological and spatial arrangement of connections in connectomes is thought to shape the segregation and integration of information and, ultimately, their computational capacity [37]. To investigate inter-species differences in the topological and spatial distribution of connections, we stratify edges into different classes in four commonly studied partitions. Partitions include: inter- and intra-modular connections (Fig. 5a), inter- and intra-hemispheric connections (Fig. 5b), connection length distribution (short, medium and long-range connections; Fig. 5c and Fig. S11), and rich-club (rich-club, feeder and peripheral connections; Fig. 5d). Overall, we find that, along the four partitions, the relative proportions of each connection class are conserved across taxonomic orders, despite differences in connection density. Collectively, this is consistent with the results from the previous sections showing that global architectural features of connectomes are consistent across phylogeny.

**FIG. 5.**
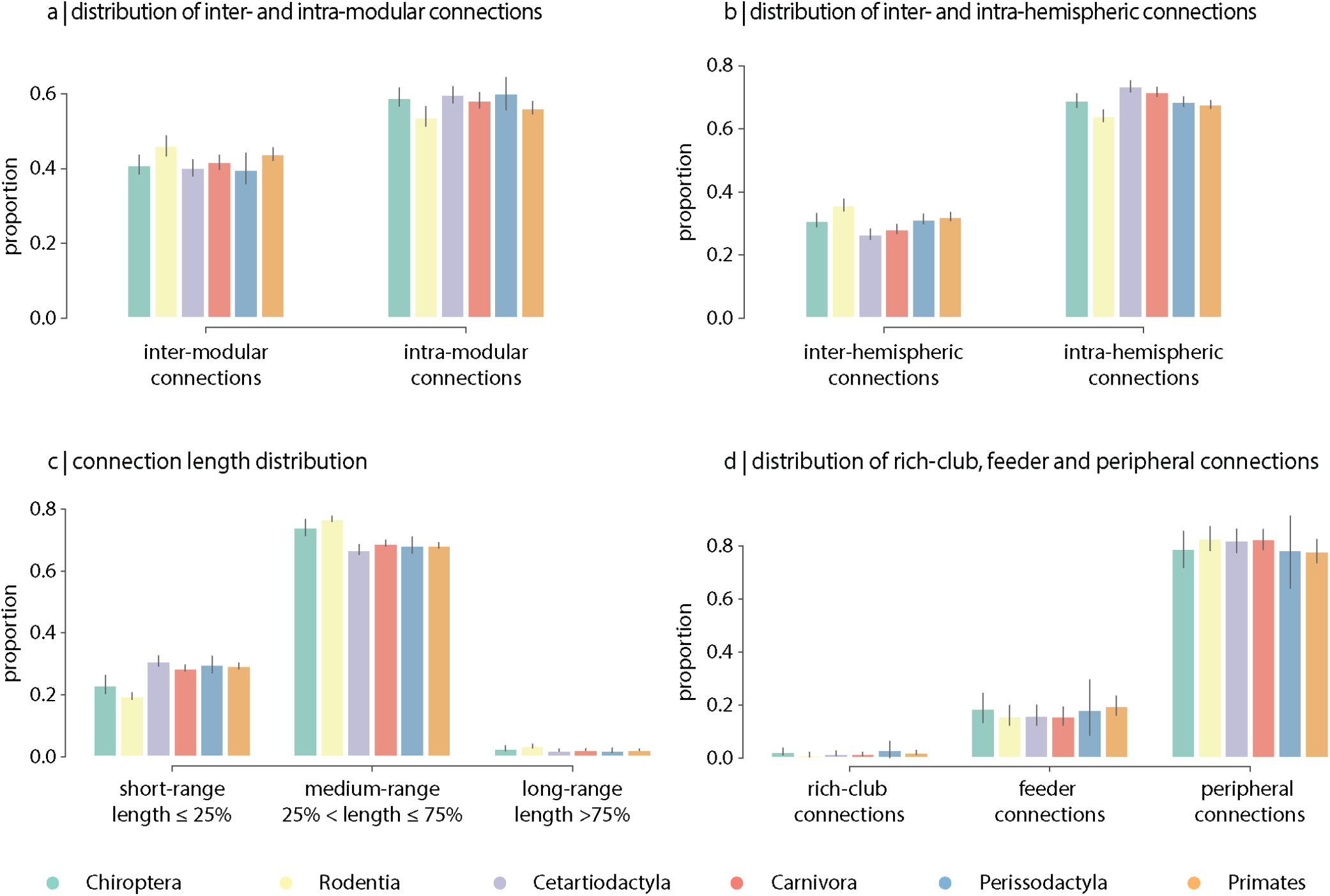
Contribution of edge types. (a) Connection density distribution for each taxononomic order. (b) Mean proportion of intra- and inter-hemispheric connections. (c) Mean proportion of short (length ≤ 25%), medium (25% < length ≤ 75%) and long connections (length ≥ 75%). (d) Mean proportion of rich-club (connecting two rich club nodes), feeder (connecting one rich club and one non-rich club node) and peripheral (connecting two non-rich club nodes) connections. Error bars indicate 95% confidence intervals.

## DISCUSSION

In the present study we chart the organization of whole-brain neural circuits across 124 mammalian species and 5 superorders. We find that connectome organization recapitulates traditional taxonomies. While all connectomes retain hallmark global features and relative proportions of edge classes, inter-species variation is driven by local regional connection profiles.

Conventional mammalian taxonomies are delineated based on the concept that a species is a group of organisms that can reproduce naturally with one another and create fertile offspring [56]. As a result, classical taxonomies based on animal morphology have largely been reconciled with emerging evidence from wholegenome sequencing [6]; namely, organisms with similar genomes display similar physical characteristics and behaviour. Our work shows that inter-species similarity – as defined by morphology, behaviour and genetics – is concomitant with the organization of neural circuits. Specifically, species that are part of the same taxonomic order tend to display similar connectome architecture, suggesting that brain network organization is under selection pressure, analogous to size, weight or color.

Which network features drive differences across taxonomic orders? Interestingly, all connectomes display consistent global hallmarks that were previously documented in tract-tracing studies, including high clustering and near-minimal path length characteristic of smallworld organization, as well as segregated network communities and densely-interconnected hub nodes [95]. The conservation in global wiring and organizational principles is further supported by a reduced difference between intra- and inter-order topological distances estimated exclusively from global features, compared to the case in which only local features are considered. Thus, relative differences between connectomes across taxonomic orders are mainly driven by local regional features. These results are in line with the idea that a brain region’s functional fingerprint — the specific computation or function that it performs by virtue of its unique firing patterns and dynamics — is determined by its underlying cortico-cortical connectional fingerprint [59, 61, 62, 76]. Accordingly, inter-species differences in functional and behavioural repertoire are likely supported by changes in local connectivity patterns. Along the same lines, our results are also consistent with the notion that neural circuit evolution involves local circuit modifications to adapt to specific challenges [10], such as extreme environmental pressures [36, 75, 84], or to support specific behaviours, such as courtship [9, 34, 35, 47, 58, 73, 82, 107], social bonding [45, 46, 53, 103], or foraging [74, 99]. How computations and cognitive functions emerge from these species-specific circuit modifications remains a key question in the field [22, 90].

It is noteworthy that the relative proportion of edge classes (inter-vs intra-modular, inter-vs intra-hemispheric, short-vs medium-vs long-range and rich-club vs feeder vs peripheral) are preserved across species. This result is reminiscent of recent work on allometric scaling that investigates how white matter connectivity scales with brain size [23]. For example, species with fewer commissural inter-hemispheric connections exhibit lower hemispheric mean-shortest-path (i.e., stronger intra-hemispheric connectivity), suggesting a similar connectivity conservation principle [3]. Likewise, using diffusion-weighted MRI data across 14 different primate species, another study reported negative allometric scaling of cortical surface area with white matter volume and corpus callosum cross-sectional area [2]. This scaling results in less space for white matter connectivity with increasing brain size, translating into larger brains with a relatively higher proportion of shortrange connections than long-range connections when compared with smaller brains [2]. Collectively these studies highlight that the architecture of neural circuits and their physical embedding are intertwined, and the distribution of connections is such that it retains consistent global architectural features across phylogeny.

The present results contribute to the emerging field of comparative connectomics [10, 59, 60, 91, 95]. Adopting a harmonized imaging protocol in a large number of mammalian species facilitates a rigorous quantitative comparison of neural circuits. Central to this are network analytic methods that map connectomes to a common space, and quantify similarities across local and global levels of organization [16, 31, 32, 59, 61, 100]. By comprehensively charting taxonomies of connectome architectures we may uncover the principles that govern the wiring of neural circuits [4]. In particular, quantitative analysis of connectome architecture across phylogeny may help to link genomics and behaviour [66]. Traditionally, taxonomic groups were defined in terms of physical morphology and behavioural repertoire [24], but these are now understood to be driven by speciation events in the genome [1, 28, 67]. However, we do not yet understand how genes influence neural circuit architecture, which in turn shapes the behavioural repertoire of an organism. By understanding how neural circuits change over phylogeny we can fill this gap and forge a link from genes to circuits to behaviour. Ultimately, the confluence of genomics, connectomics and behaviour may help to triangulate towards a more well-rounded view of speciation [42], and their simultaneous investigation can further illuminate the link between structure and function in brain networks [14, 88, 89].

The present work must be considered with respect to multiple limitations. First, uniformly parcellating brains of different size into the same number of nodes facilitates comparison of network architecture, but potentially obscures biologically important regional differences. Second, many species are represented by a single individual. Although we focus on orders rather than individual species and there is high within-species reliability [3], the analyses do not capture individual variability within species. Third, all connectomes are reconstructed using diffusion weighted imaging, which is subject to both systematic false positives and false negatives [54, 80]. While a uniform, high-resolution *ex vivo* scanning protocol allows for systematic comparisons among species, and our results recapitulate findings from tract tracing studies, future work in comparative connectomics will benefit from technological and analytical advances in neural circuit mapping. Fourth, evolutionary circuit modifications may not occur at the level of large-scale white matter, but at finer scales involving smaller nuclei or physiological events not accessible by diffusion imaging, such as up- or down-regulation of neurotransmitter receptors [10]. Nevertheless, the strikingly consistent taxonomic and phylogenetic relationships revealed by connectome analysis remain, and suggest that macroscale connectivity, as measured by diffusion MRI, is informative of species similarities and differences across taxonomic orders.

By encoding connectomes into a common frame of reference, we quantitatively assess neural circuit architecture across the mammalian phylogeny. We find that connectome organization recapitulates previously established taxonomic relationships. Collectively, these findings set the stage for future mechanistic studies to trace the link between genes to neural circuits and ultimately to behaviour, and offer new opportunities to explore how changes in brain network structure across phylogeny translate into changes in function and behavior.

## METHODS

### Brain samples

The mammalian MRI (MaMI) database includes a total of 225 ex vivo diffusion and T2- and T1-weighted brain scans of 125 different animal species (Supplementary Table 1). No animals were deliberately euthanized for the present study. All brains were collected based on incidental death of animals in zoos in Israel or natural death collected abroad, and with the permission of the national park authority (approval no. 2012/38645) or its equivalent in the relevant countries. All scans were performed on excised and fixated tissue. Animals’ brains were extracted within 24 h of death and placed in formaldehyde (10 percent) for a fixation period of a few days to a few weeks (depending on the brain size). Approximately 24 h before the MRI scanning session, the brains were placed in phosphate-buffered saline for rehydration. Given the limited size of the bore, small brains were scanned using a 7-T 30/70 Biospec Avance Bruker system, whereas larger brains were scanned using a 3-T Siemens Prisma system. To minimize image artifacts caused by magnet susceptibility effects, the brains were immersed in fluorinated oil (Flourinert, 3M) inside a plastic bag during the MRI scanning session.

### MRI acquisition

A unified MRI protocol was implemented for all species. The protocol included high-resolution anatomical scans (T2- or T1-weighted MRI), which were used as an anatomical reference, and diffusion MR scans. Diffusion MRI data were acquired using high angular resolution diffusion imaging (HARDI), which consists of a series of diffusion-weighted, spin-echo, echo-planar-imaging images covering the whole brain, scanned in either 60 (in the 7-T scanner) or 64 gradient directions (in the 3-T scanner) with an additional 3 non-diffusion-weighted images (B0). The b value was 1,000 smm^−2^ in all scans. In the 7-T scans, TR was longer than 12,000 ms (depending on the number of slices), TE was 20 ms and Δ/*δ* = 10/4.5 ms. In the 3-T scans, TR was 3,500 ms, with a TE of 47 ms and Δ/*δ* = 17/23 ms.

To linarly scale according to brain size the twodimensional image pixel resolution (per slice), the size of the matrix remained constant across all species (128 x 96). Due to differences in brain shape, the number of slices varied between 46 and 68. Likewise, the number of scan repetitions and the acquisition time were different for each species, depending on brain size and desired signal to noise ratio (SNR) levels. To keep SNR levels above 20, an acquisition time of 48 h was used for small brains (~ 0.15ml), and 25 min for large brains (>1,000 ml). SNR was defined as the ratio of mean signal strength to the standard deviation of the noise (an area in the nonbrain part of the image). Full details are provided in Assaf et al. [3].

### Connectome reconstruction

The ExploreDTI software was used for diffusion analysis and tractography [50]. The following steps were used to reconstruct fiber tracts:

1. To reduce noise and smooth the data, anisotropic smoothing with a 3-pixel Gaussian kernel was applied.
2. Motion, susceptibility and eddy current distortions were corrected in the native space of the HARDI acquisition.
3. A spherical harmonic deconvolution approach was used to generate fiber-orientation density functions per pixel [93], yielding multiple (n>= 1) fiber orientations per voxel. Spherical harmonics of up to fourth order were used.
4. Whole-brain tractography was performed using a constrained spherical deconvolution (CSD) seed point threshold similar for all samples (0.2) and a step length half the pixel size.

The end result of the tractography analysis is a list of streamlines starting and ending between pairs of voxels. Recent studies have shown that fiber tracking tends to present a bias where the vast majority of end-points reside in the white matter [93]. To avoid this, the CSD tracking implemented here insures that approximately 90% of the end-points reside in the cortical and subcortical grey matter.

### Network generation and analysis

Before the reconstruction of the networks, certain fiber tracts were removed from the final list of tracts. These include: external projection fibers that pass through the cerebral peduncle, as well as cerebellar connections. Inner-hemispheric projections, such as the thalamic radiation, were included in the analysis. Brains were parcellated into 200 nodes using a *k*-means clustering algorithm. All the fiber end-point positions were used as input, and cluster assignment was done based on the similarity in connectivity profile between pairs of end-points. Therefore, vertices with similar connectivity profile have a higher chance of grouping together. The clustering was performed twice, once for each hemisphere. Nodes were defined as the mass center of the resulting 200 clusters. Connectivity matrices were generated by indexing the number of streamlines between any two nodes [3]. We note that, even though the use of a non-anatomically informed parcellation does not allow for direct comparison of individual regions across species in terms of anatomical location, it does ensure that the resulting connectivity matrices include the same number of regions across species.

### Controlling for the scanning resolution and acquisition parameters

As the size of the matrix was kept constant across species (i.e., 200 nodes), voxel dimensions were linearly scaled with brain volume, thus resulting in different scanning resolutions across the samples. To verify that this was a reasonable assumption, several tests were performed: (1) The diffusion-based connectome of the mouse was previously compared against one derived from tract-tracing (see [3] for details), obtaining a strong correlation between both networks. (2) Results on the connectivity conservation principle presented in [3] were invariant to different scanning and parcellation parameters across nine different species. (3) The diffusion weighted imaging method was able to reconstruct specific ground truth fiber systems across brains, and these fiber bundles scaled in size with brain volume.

### Spectral distance

To estimate similarities between species’ connectome organization, we computed the Laplacian eigenspectrum of each graph. The Laplacian eigenspectrum acts like a “radiography” of the graph and summarizes distinct aspects of the underlying topology [7, 8, 30, 40, 41, 70]. We considered the normalized Laplacian matrix *L* for undirected graphs with (symmetric) adjacency matrix *A. L* is defined as *L* = *I* – *D*^−1^ *A*, where *D* is a diagonal matrix with *D*(*i*, *i*) = deg *i*, and deg *i* is the binary degree of vertex *i*. Specifically,

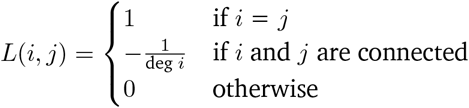

with *i* and *j* representing two vertices of the graph. The Laplacian spectrum is then given by the set of all the eigenvalues of *L*. Importantly, the eigenspectrum of the normalized Laplacian has the advantage that all eigenvalues are in the range [0, 2] [26], facilitating comparison across species. Furthermore, the normalized Laplacian is unitarily equivalent to the symmetric normalized Laplacian [26], i.e., 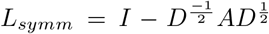, thus the eigenvalues of both Laplacians are real.

We smoothed the Laplacian eigenvalue distribution (i.e., *λ*_1_, *λ*_2_,…, *λ_n_*) by convolving eigenvalue frequencies with a Gaussian kernel. The new kernel density estimation is given by:

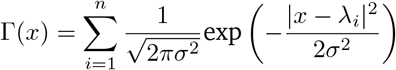

with *n* being the number of eigenvalues in the approximated distribution, and *σ* being a smoothing factor of 0.015. We used a step of 0.001, which resulted in a total of *n* = 2000 eigenvalues. The approximated distribution was normalized such that area under the curve is one. The spectral distance between every pair of animal species was estimated as 1 minus the cosine similarity of their smoothed (normalized) Laplacian eigenvalue distributions, where eigenvalue distributions were assumed to be vectors in a high dimensional space.

### Topological distance

An alternative way to estimate inter-species distances in connectome organization is to compute the correlation between their network features. We estimated a set of local and global graph theory measures of the connectivity matrix. Local measures include node degree, clustering coefficient, node betweenness and closeness. Global measures include characteristic path length, transitivity and assortativity. We included both the binary and the weighted versions of these measures. We constructed a vector of local and global topological features for every animal species. Because there are as many local features as nodes in a network, we only used the average and the standard deviation of these measures. Similar to the spectral distance, the topological distance between every pair of animal species was estimated as 1 minus the cosine similarity of their topological feature vectors. All local and global features were estimated using the Python version of the Brain Connectivity Toolbox (https://github.com/aestrivex/bctpy; [77]). Definitions of these topological metrics can be found below.

#### Local features

- *Degree* (*bin*). Number of connections that a node participates in:

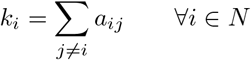

where *a_ij_* = 1 if nodes *i* and *j* are connected, otherwise *a_ij_* = 0. *N* corresponds to the set of all nodes in the graph [77].
- *Degree* (*wei*). Sum of connection weights that a node participates in:

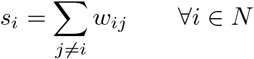

where *w_ij_* corresponds to the connection weight between nodes *i* and *j*. *N* corresponds to the set of all nodes in the graph [77].
- *Clustering* (*bin*). Proportion of transitive closures (closed triangles) around a node, i.e., the fraction of neighbor nodes that are neighbors of each other:

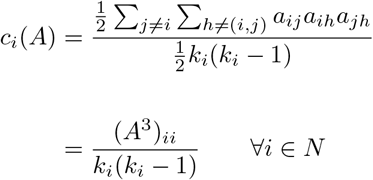

where *A* corresponds to the binary adjacency matrix of the graph, (*A*^3^)_*ii*_ is the *i*-th element of the main diagonal of *A*^3^ = *A* · *A* · *A*, and *k_i_* is the degree of node *i* [77, 101].
- *Clustering* (*wei*). Mean “intensity” of triangles around a node:

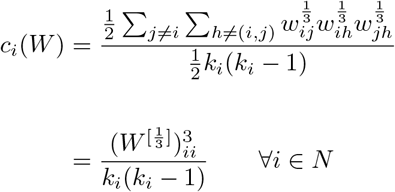

where *W* corresponds to the weighted connectivity matrix, 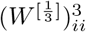 is the *i*-th element of the main diagonal of 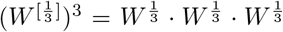, and *k_i_* is the degree of node *i* [72, 77].
- *Shortest path length* (*bin*). Minimum geodesic distance between pairs of nodes:

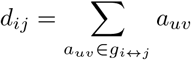

where *g*_*i*↔*j*_ is the shortest geodesic path between nodes *i* and *j*. Note that *d_ij_* = ∞for all disconnected pairs *i*, *j* [77].
- *Shortest path length* (*wei*). Minimum weighted distance between pairs of nodes:

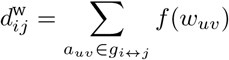

where *f* is a map (e.g., the inverse) from weight to length and *g*_*i*↔*j*_ is the shortest weighted path between nodes *i* and *j* [77].
- *Betweenness* (*bin*). Proportion of shortest (geodesic) paths in the graph that traverse a node:

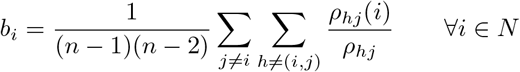

where *ρ_hj_* is the number of shortest (geodesic) paths between nodes *h* and *j*, *ρ_hj_*(*i*) is the number of shortest (geodesic) paths between nodes *h* and *j* that pass through *i*, and *n* is the number of nodes in the graph [21, 39, 48, 77].
- *Betweenness* (*wei*). Same as *betweenness* (*bin*), but shortest paths are estimated on the respective weighted graph [21, 39, 48, 77].
- *Closeness* (*bin*). Mean shortest (geodesic) path length from a node to all other nodes in the network:

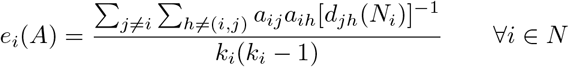

where *d_jh_*(*N_i_*) is the shortest (geodesic) path length between nodes *j* and *h*, that contains only neighbors of *i*, and is estimated on the corresponding binary adjacency matrix *A* [49, 77].
- *Closeness* (*wei*). Mean shortest (weighted) path from a node to all other nodes in the network:

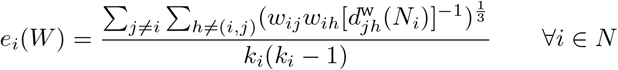

where 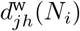 is the length of the shortest (weighted) path between nodes *j* and *h*, that contains only neighbors of *i*, and is estimated on the corresponding weighted adjacency matrix *W* [49, 77].

#### Global features

- *Characteristic path length* (*bin*). Average shortest (geodesic) path length between all pairs of nodes in the graph:

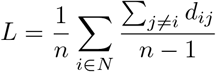

where *d_ij_* is the shortest (geodesic) path length between nodes *i* and *j*, and *n* is the number of nodes in the graph [77, 101].
- *Characteristic path length* (*wei*). Average shortest (weighted) path length between all pairs of nodes in the graph:

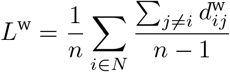

where 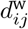 is the shortest (weighted) path length between nodes *i* and *j*, and *n* is the number of nodes in the graph [77, 101].
- *Transitivity* (*bin*). Ratio between the observed number of closed triangles and the maximum possible number of closed triangles:

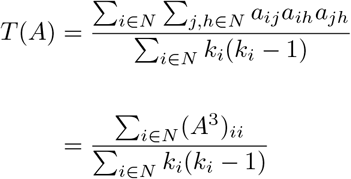

where *A* is the binary adjacency matrix, and *k_i_* is the degree of node *i* [69, 77].
- *Transitivity* (*wei*). Ratio between the “intensity” of observed closed triangles and the maximum possible “intensity” of closed triangles:

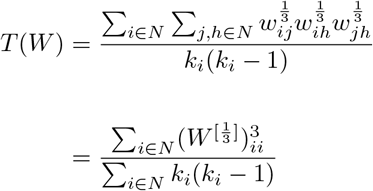

where *W* is the weighted connectivity matrix, and *k_i_* is the degree of node *i* [69, 77].
- *Assortativity* (*bin*). Correlation coefficient between the degree of a node and the mean degree of its neighbours:

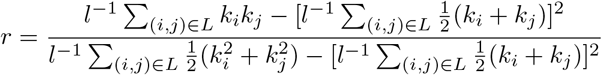

where *l*^−1^ is the inverse of the number of links in the graph, *k_i_* is the degree of node *i*, and *L* is the set of links in the graph [68, 77].
- *Assortativity* (*wei*). Correlation coefficient between the weighted degree of a node and the mean wighted degree of its neighbour:

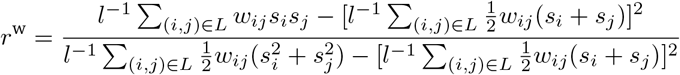

where *l*^−1^ is the inverse of the number of links in the graph, *s_i_* is the weighted degree of node *i*, and *L* is the set of links in the graph [51, 77].

### Small-world organization

We use the index proposed in [44] to measure connectomes’ small-worldness level. The index is given by:

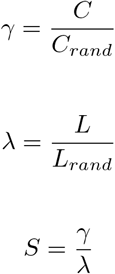

where *L* and *C* are the corresponding characteristic path length and clustering coefficient of each connectome, respectively, and *L_rand_* and *C_rand_* are the corresponding average quantities for a set of 1000 randomly rewired graphs that preserve the degree sequence and distribution of the nodes [64]. A network is said to possess a small world architecture if *S* ≥ 1, that is, if it situates above the identity line in a *γ* vs *λ* plot.

### Community detection

We used the Louvain algorithm to determine the optimal community structure of connectomes [19]. Briefly, this algorithm extracts communities from large networks by optimizing a modularity score. Here we use the Q-metric as the objective function [19, 38]:

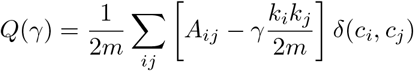

where *A* corresponds to the adjacency matrix of the network, *k_i_* is the degree of node *i*, *m* is the sum of all connections in the graph, *c_i_* is the community affiliation of node *i*, and *δ* is the Kronecker delta function (i.e., *δ*(*x*, *y*) = 1 if *x* = *y*, 0 otherwise). The size of the partition is controlled by a resolution parameter *γ* (higher *γ* values result in a larger number of modules). Because the Louvain method is a greedy algorithm, we first found multiple (250) optimal partitions at *γ* = 1, and then we determined a single partition using a consensus clustering approach [15].

### Classification of edges

Connectomes’s connections were classified into different categories based on four criteria. The first criterion is based on the modular structure of the network and classifies connections depending on whether they link brain regions within the same module (i.e., intra-modular) or regions across different modules (i.e., inter-modular); the second criterion is whether connections link brain regions within the same hemisphere (i.e., intra-hemispheric connections) or across hemispheres (inter-hemispheric connections); the third criterion is based on the physical length of the connections (i.e., short-, medium- and long-range connections), and the fourth criterion is based on the rich-club structure of the network (i.e., rich-club, feeder and peripheral connections).

#### Inter-vs intra-modular connections

To classify connections as being either inter- or intra-modular, a consensus clustering algorithm was applied on each connectome to determine a partition of the network into different modules (see *Community detection*). Once modules are identified, inter-modular connections correspond to those linking brain regions across different modules, whereas intra-modular connections correspond to those linking brain regions belonging to the same module.

#### Connection length

Euclidean distance between regions’ centers was used as a proxy for connection length. To subdivide connections into short-, medium- and long-range, connection lengths were estimated as a percentage with respect to the maximum distance between regions. Short-range connections correspond to those that are less or equal to 25%of the maximum distance; medium-range connections are above 25%but less than or equal to 75%of the maximum distance; and long-range connections are those above 75%of the maximum distance.

#### Rich-club vs feeder vs peripheral connections

To classify edges as being either rich-club, feeder or peripheral connections, it is necessary to identify first the rich-club of hubs in the network, that is, the densely interconnected core of nodes that have a disproportionately high number of connections [96, 98]. To do so, we compute the rich-club coefficients Φ(*k*) across a range of degree *k* of the unweighted (binary) connectomes. For binary networks, all nodes that show a degree ≤ *k* are removed from the network, and for the remaining set of nodes (i.e., a sub-graph), the rich club coefficient is estimated as the ratio of connections present in the sub-graph, to the total number of possible connections that would be present if the resulting sub-graph was fully connected. Formally, the rich-club coefficient is given by [27, 65, 109]:

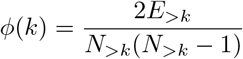

In random networks, such as the Erdős-Rényi model, nodes with a higher degree have a higher probability of being interconnected by chance alone, thus showing an increasing function of Φ(*k*). For this reason, the rich-club coefficient is typically normalized relative to a set of *m* comparable random networks of equal size and node degree sequence and distribution [27, 64, 65]. The normalized rich-club coefficient is then given by:

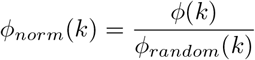

where *ϕ_random_* corresponds to the average rich-club coefficient over the *m* random networks. In our particular case *m* = 1000. An increasing normalized coefficient Φ_*norm*_ > 1 over a range of *k* reflects the existence of rich-club organization.

To assess the statistical significance of rich-club organization we used permutation testing [13, 97]. Briefly, the population of the *m* random networks yields a null distribution of rich-club coefficients. For the range of *k* expressing rich-club organization (i.e., Φ*_norm_* > 1), we tested whether *ϕ*(*k*) significantly exceeds Φ*_random_*(*k*).Next, we identify the *k*^th^ level at which the maximum significant Φ_*norm*_ occurs. Nodes with a degree ≥ *k* are said to belong to the *k*^th^-core of the network. Next, we identify the hubs, that is, nodes whose degree is above the average degree of the network plus one standard deviation. Therefore, rich-club nodes were identified as those nodes that are both hubs and belong to the *k*^th^-core of the network.

Once rich-club nodes are identified, rich-club connections are defined as edges between rich-club nodes; feeder connections are edges connecting rich-club to non-rich-club nodes, and peripheral connections are edges between non-rich-club nodes.

### Controlling for replicas

Because some of the species have multiple scans, this could bias the distribution of intra- and inter-order distances, which could be dominated by those species with a large number of replicas. To account for that, we randomly sample a single connectome per species, and we calculated inter-species distances. We repated this procedure iteratively 10,000 times. The reported intra- and inter-order distance distributions correspond to the average distances across iterations.

### Controlling for density

Some of the graph features used for the estimation of the topological distance are highly dependant on the density of the network. To regress out the effects of network density on a graph feature, a univariate linear and an exponential model are fitted using density as the explanatory variable, and each feature as response variable. That is:

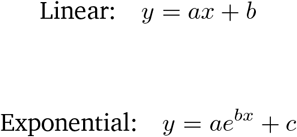

where *y* represents network features and *x* corresponds to density. Network features are then replaced by the residuals of the model. The decision to fit either a linear, an exponential or no model at all was based on the variance explained by the model or *R*^2^. The model with the largest *R*^2^ is selected. Only those features with *R*^2^ > 0.1 were controlled to account for density (features in Fig. S9 with a regression line).

## Acknowledgments

BM acknowledges support from the Natural Sciences and Engineering Research Council of Canada (NSERC), Canadian Institutes of Health Research (CIHR), Brain Canada Foundation Future Leaders Fund, the Canada Research Chairs (CRC) Program and the Healthy Brains for Healthy Lives (HBHL) initiative. GL acknowlegdes support from NSERC, the CRC Program, and the Canada CI-FAR AI Cahir Program.

---

**TABLE S1:**
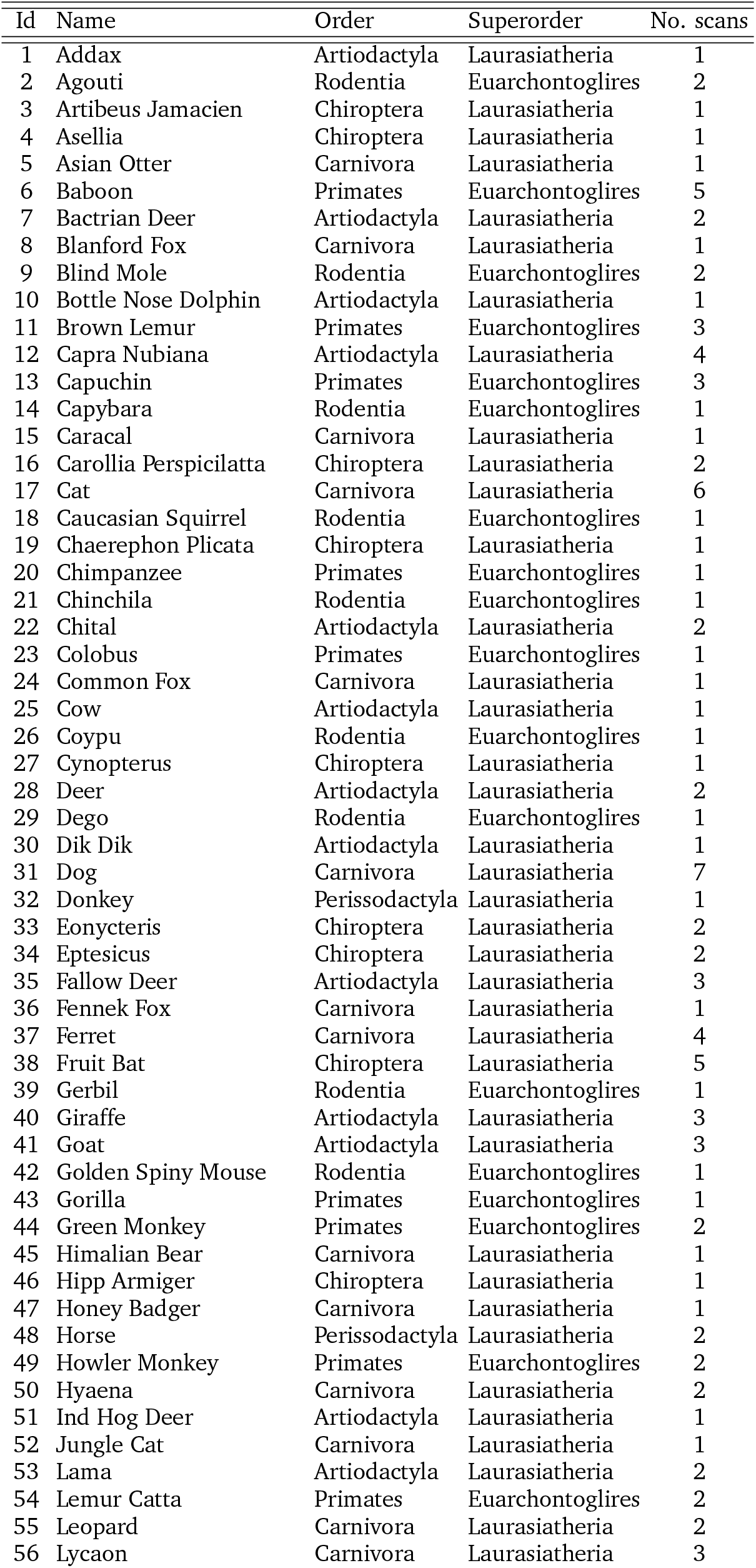
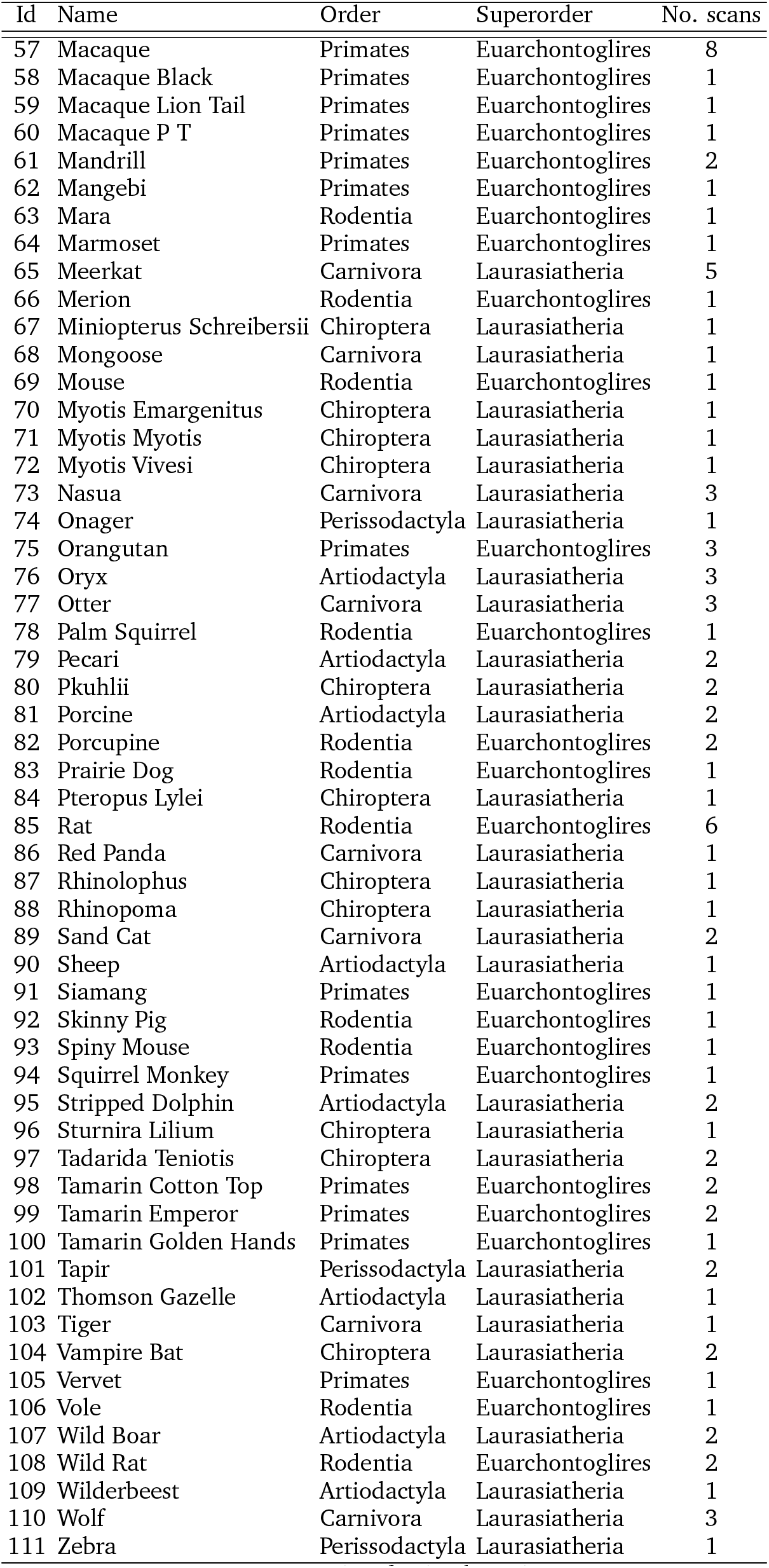
List of animal species

**FIG. S1.**
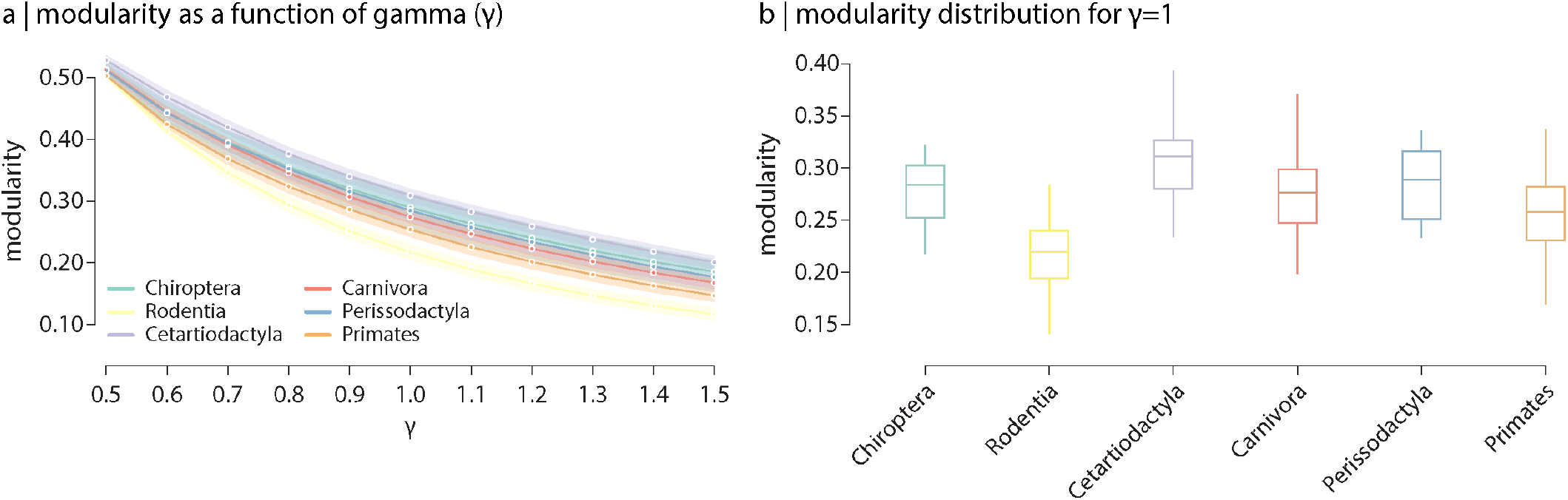
Modularity. (a) Modularity as a function of resolution the resolution parameter *γ*, which controls for the size of the identified modules. (b) Modularity distributions for each taxonomic order (*γ* = 1).

**FIG. S2.**
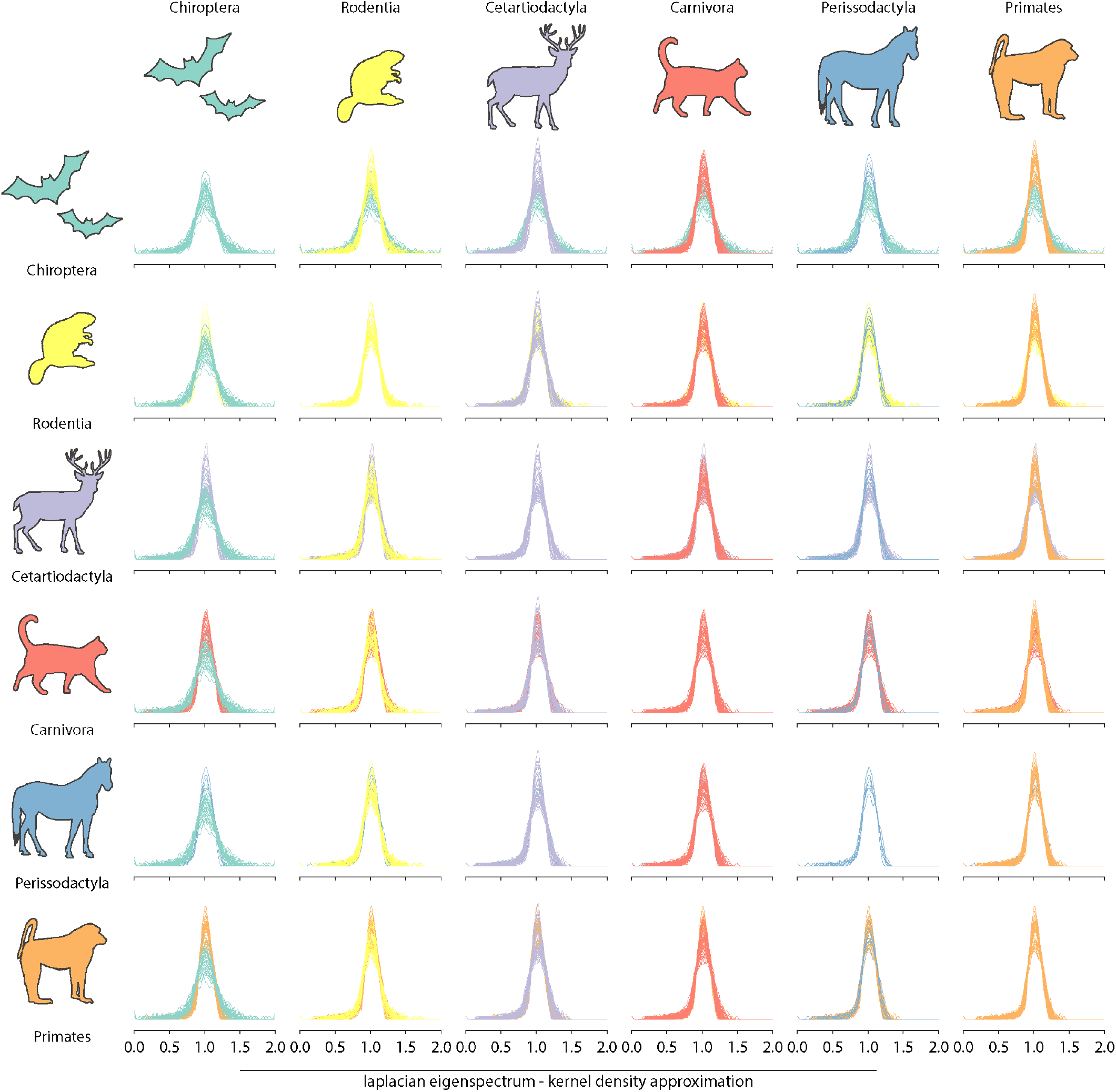
Laplacian eigenspectra. Spectral plots were obtained by convolving the eigenspectrum of the normalized Laplacian matrix of the graph with a Gaussian kernel. The eigenvalues of the normalized Laplacian of the connectivity matrix, and their multiplicities, capture distinct topological properties of the graph [7, 8, 30, 40, 41, 70], thus acting like a radiography of its underlying topology. More importantly, it has the advantage of situating graphs of different sizes and with non-homologous node correspondence in a common frame of reference in which they can be compared.

**FIG. S3.**
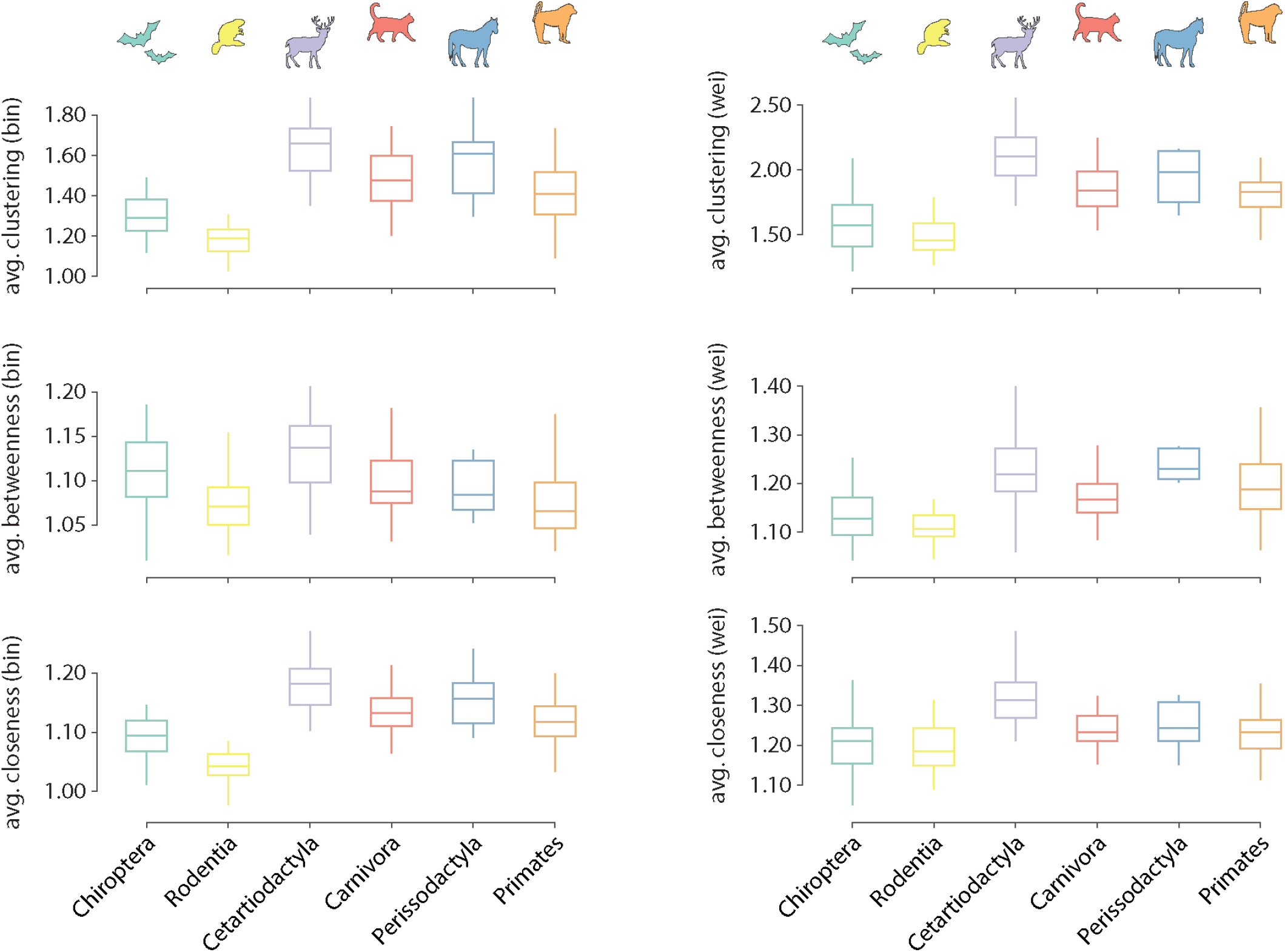
Distribution of local graph features across orders. Distributions of average local features are shown for each order. Features are normalized relative to a set of 1000 randomly rewired graphs that preserve the degree sequence and distribution of the nodes [64]. Features are computed for both the binary (left) and weighted (right) connectomes.

**FIG. S4.**
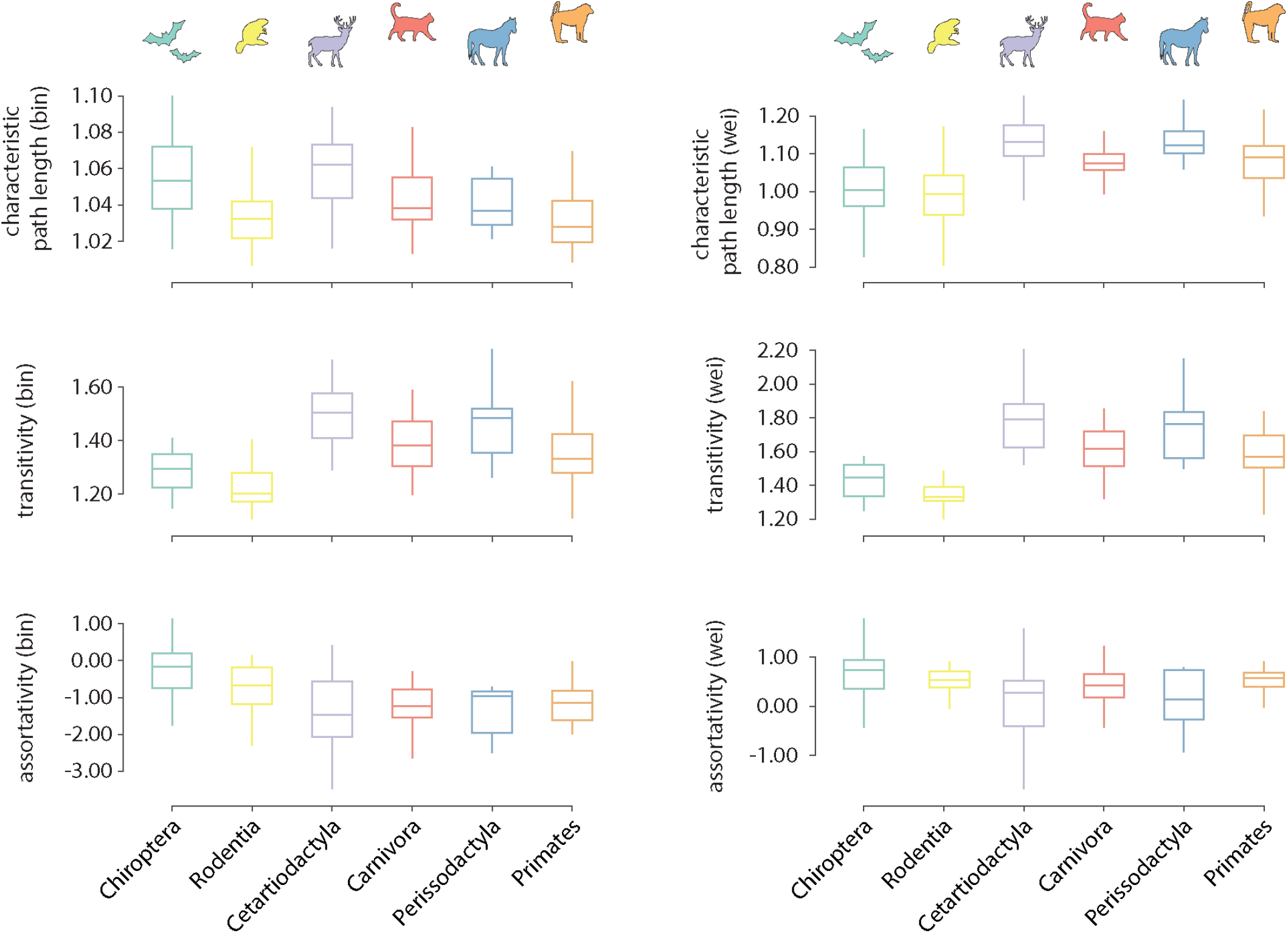
Distribution of global graph features across orders. Distributions of global features are shown for each order. Features are normalized relative to a set of 1000 randomly rewired graphs that preserve the degree sequence and distribution of the nodes [64]. Features are computed for both the binary (left) and weighted (right) connectomes.

**FIG. S5.**
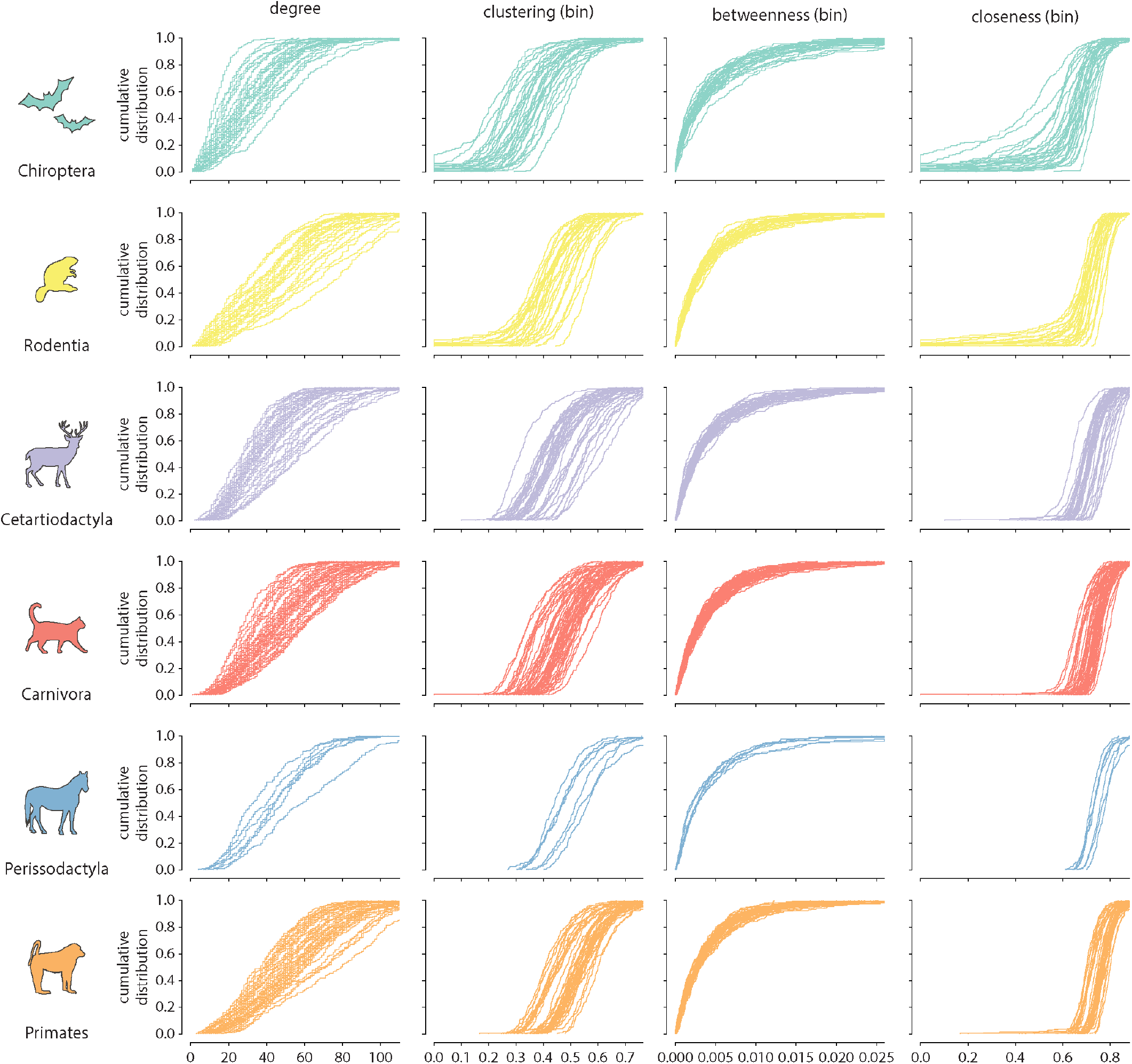
Cumulative distributions of binary local graph features across orders. The cumulative distributions of individual features are shown for each individual sample within each order.

**FIG. S6.**
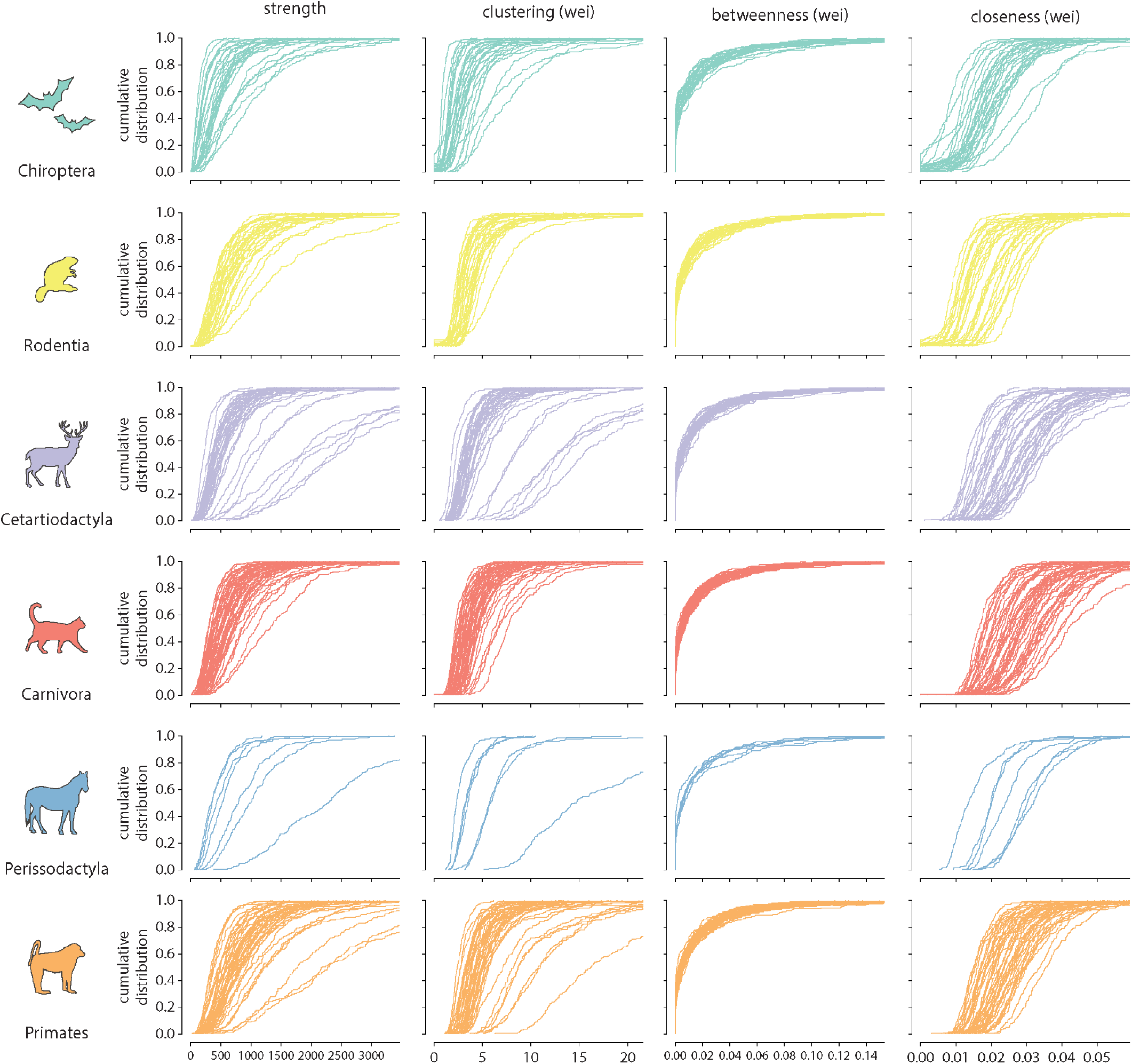
Cumulative distributions of weighted local graph features across orders. The cumulative distributions of individual features are shown for each individual sample within each order.

**FIG. S7.**
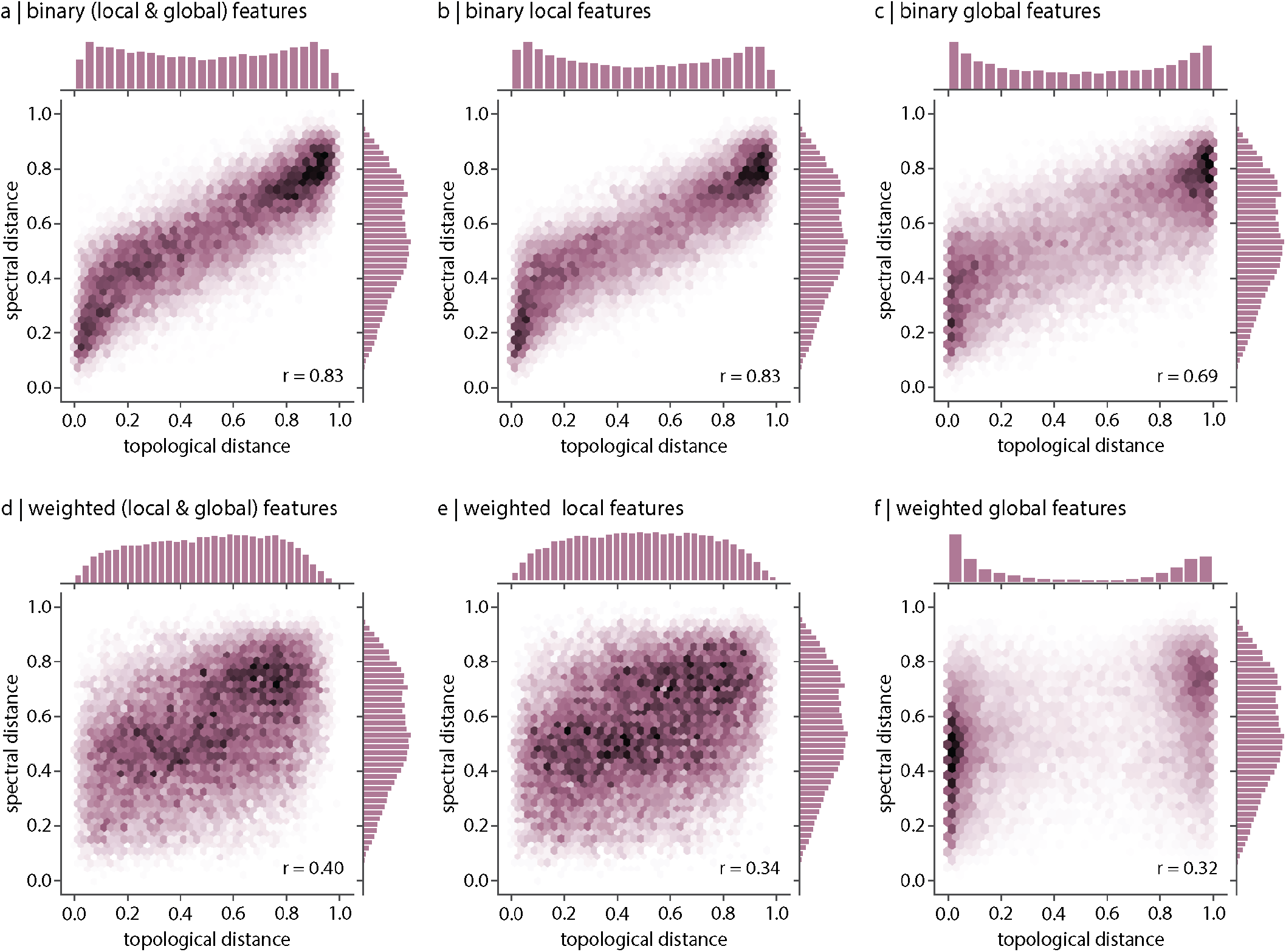
Relationship between spectral and topological distance. Correlations between inter-species distances computed using topological distance (abscissa) and spectral distance (ordinate). Correlations are shown for (a) binary local and global features, (b) binary local features, (c) binary global features, (d) weighted local and global features, (e) weighted local features and (f) weighted global features.

**FIG. S8.**
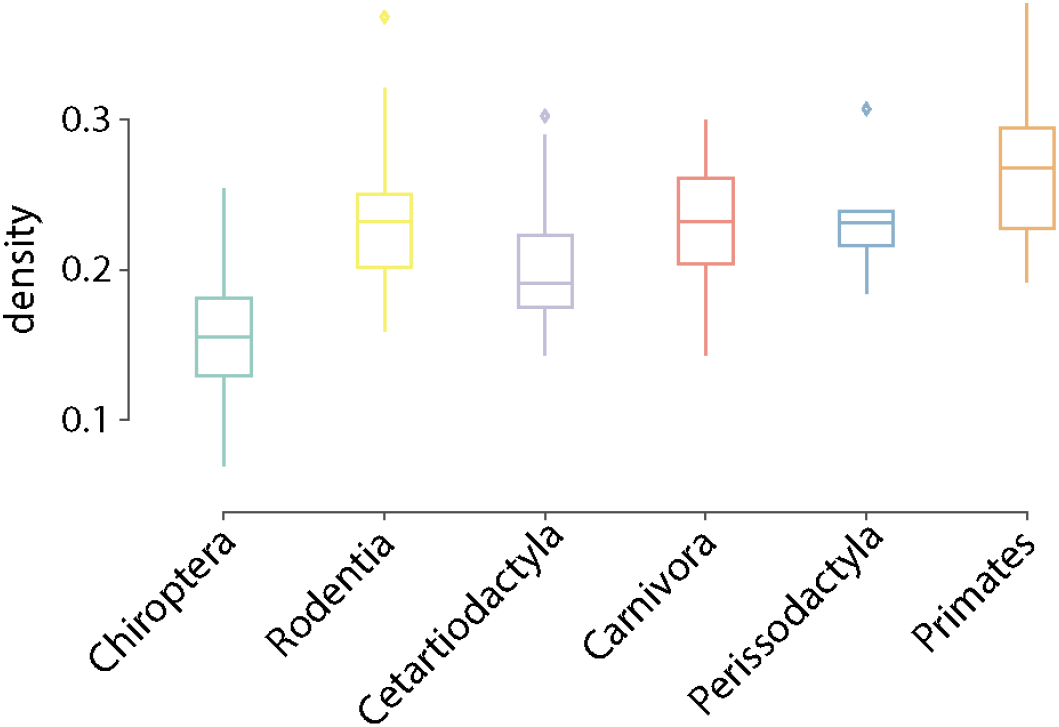
Network density. Distribution of network density is shown for each taxonomic order. Connection density is estimated as the ration of existent connections to the total number of possible connections.

**FIG. S9.**
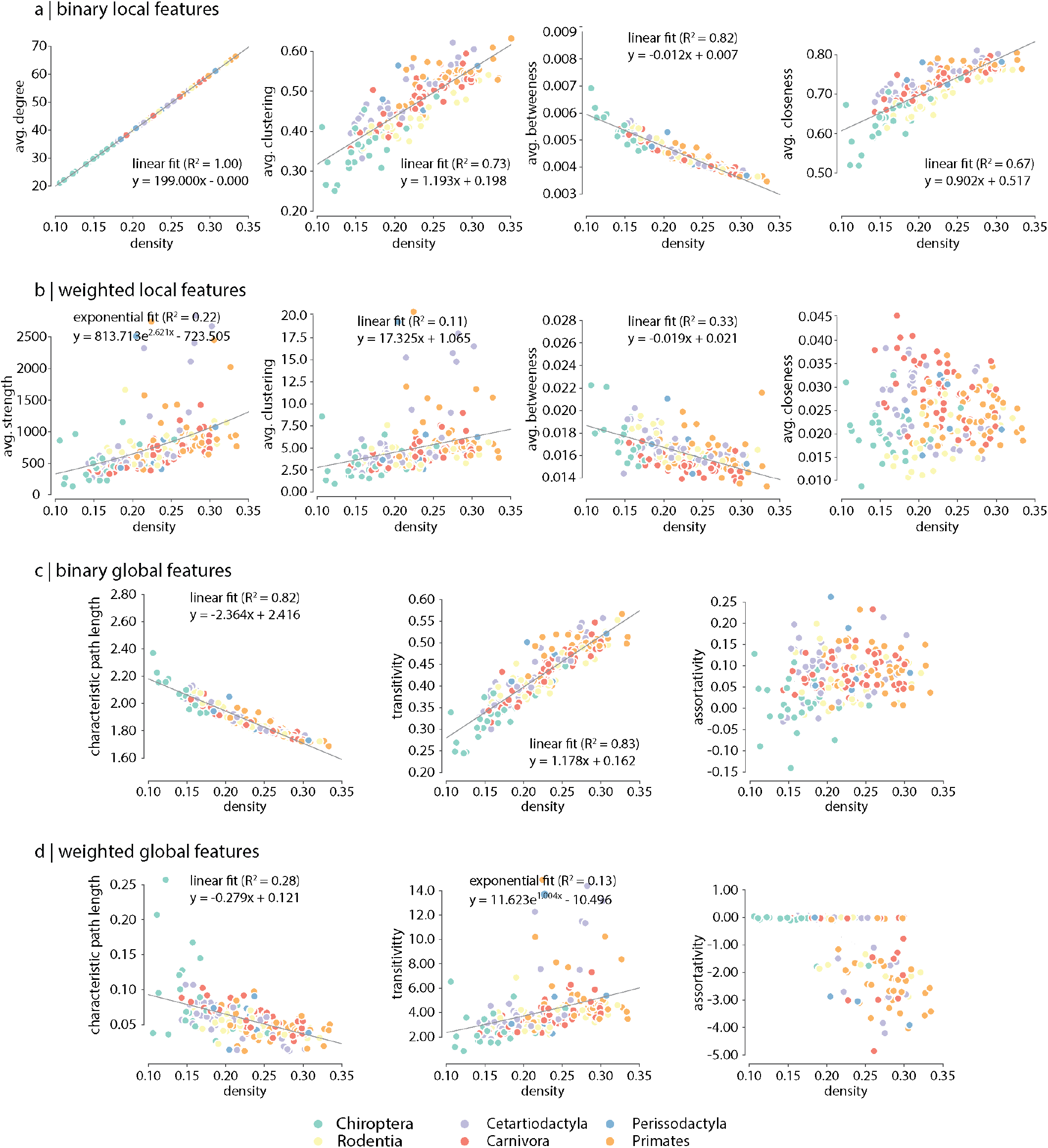
Controlling for network density. Network density is regressed out from topological features (see *Methods*).

**FIG. S10.**
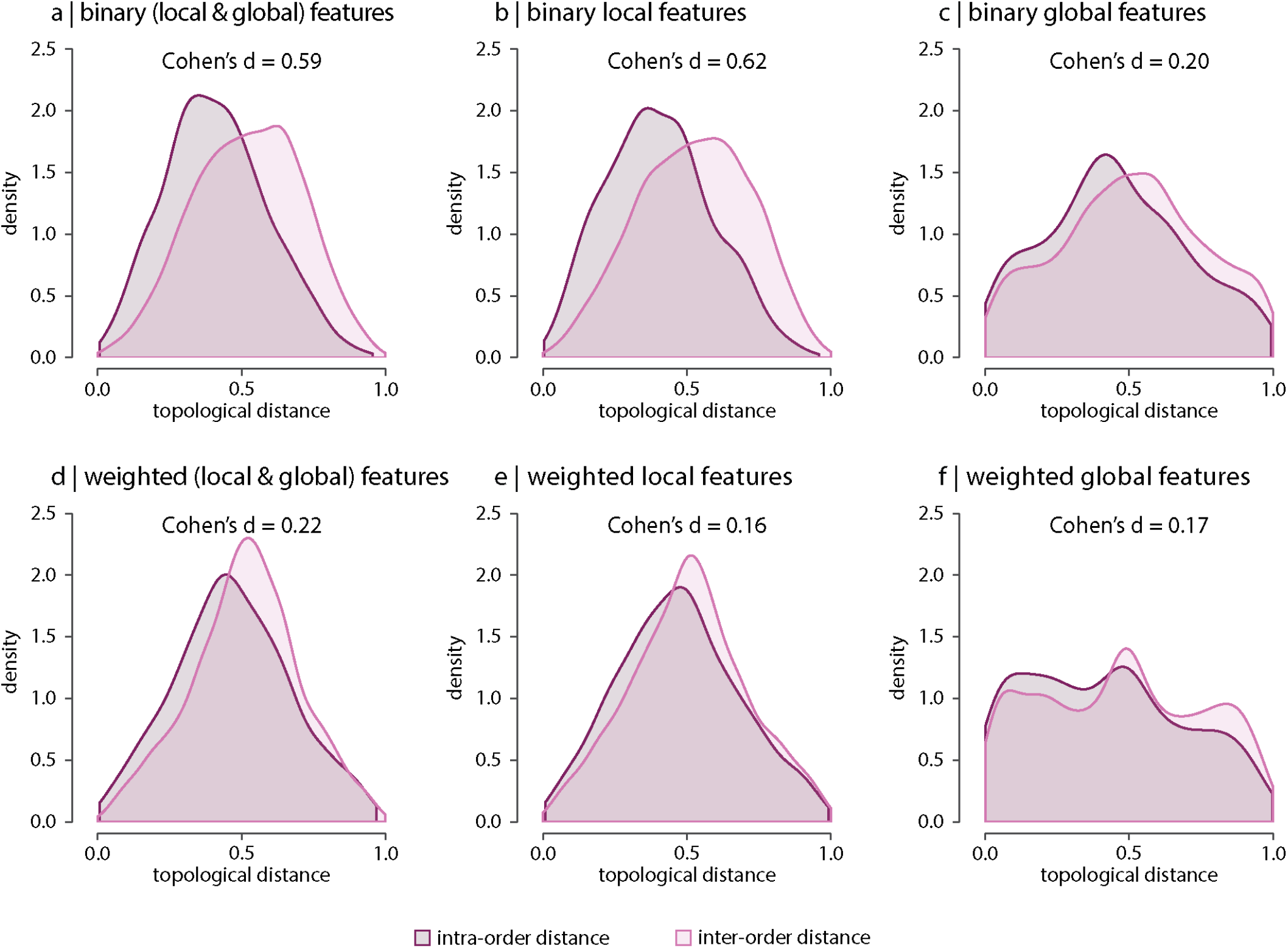
Contribution of network features after controlling for connection density. Network density is regressed out from topological features (see *Methods*), and topological distance is computed using multiple local and global connectome features. Plots show intra- and inter-order distances when using only (a) binary local and global features, (b) binary local features, (c) binary global features, (d) weighted local and global features, (e) weighted local features and (f) weighted global features after controlling for connectivity density. Local features include the average and standard deviation of degree, clustering, betweenness, and closeness. Global features include characteristic path length, assortativity and transitivity. Effect sizes correspond to the Cohen’s *d* estimator from a two-sample Welch’s t-test. Equivalent conclusions are drawn if common-language effects sizes from a two-sample Wilcoxon-Mann-Whitney rank-sum test are used. In all cases, the difference in the means and medians of intra- and inter-order distance distributions is statistically significant (*P* < 10^−4^).

**FIG. S11.**
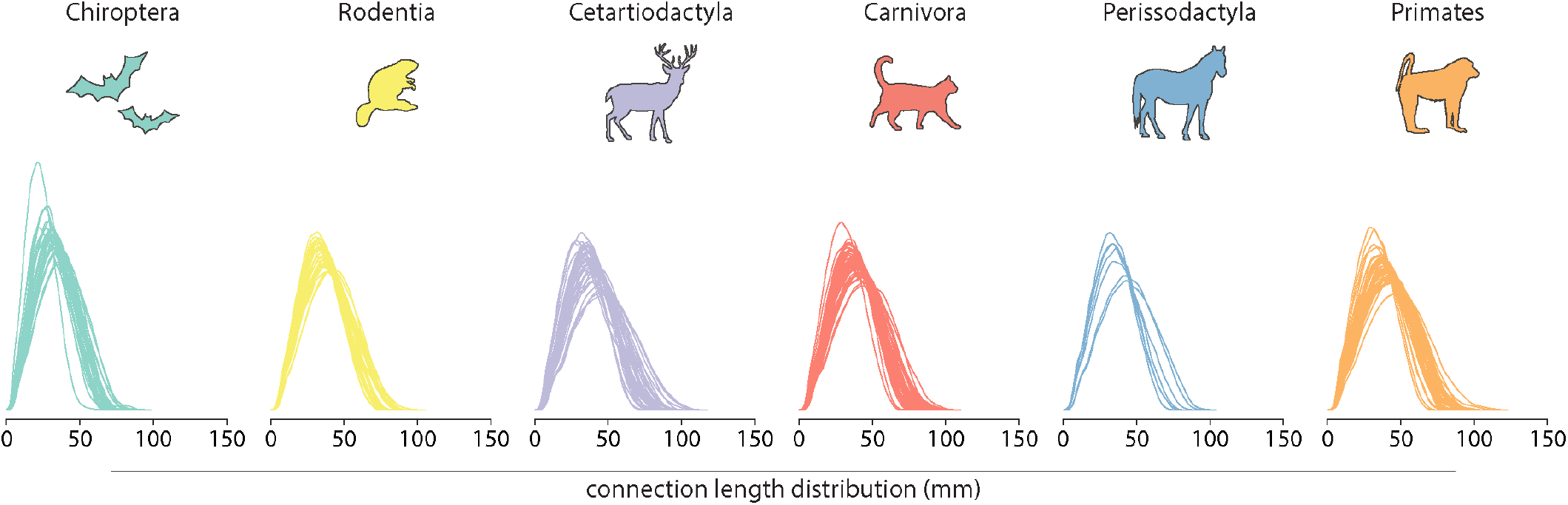
Connection length distribution. The connection length distributions of individual species for each taxonomic order.

## References

[1] Álvarez-Carretero, S., Tamuri, A. U., Battini, M., Nascimento, F. F., Carlisle, E., Asher, R. J., Yang, Z., Donoghue, P. C., and Dos Reis, M. (2021). A specieslevel timeline of mammal evolution integrating phylogenomic data. Nature, pages 1–8.

[2] Ardesch, D. J., Scholtens, L. H., de Lange, S. C., Roumazeilles, L., Khrapitchev, A. A., Preuss, T. M., Rilling, J. K., Mars, R. B., and van den Heuvel, M. P. (2021). Scaling principles of white matter connectivity in the human and nonhuman primate brain. Cereb Cortex.

[3] Assaf, Y., Bouznach, A., Zomet, O., Marom, A., and Yovel, Y. (2020). Conservation of brain connectivity and wiring across the mammalian class. Nat Neurosci, 23(7):805–808.

[4] Avena-Koenigsberger, A., Goñi, J., Solé, R., and Sporns, O. (2015). Network morphospace. J Roy Soc Interface, 12(103):20140881.

[5] Avena-Koenigsberger, A., Misic, B., and Sporns, O. (2018). Communication dynamics in complex brain networks. Nat Rev Neurosci, 19(1):17–33.

[6] Baker, R. J. and Bradley, R. D. (2006). Speciation in mammals and the genetic species concept. J Mammal, 87(4):643–662.

[7] Banerjee, A. and Jost, J. (2008). On the spectrum of the normalized graph laplacian. Linear algebra and its applications, 428(11-12):3015–3022.

[8] Banerjee, A. and Jost, J. (2009). Graph spectra as a systematic tool in computational biology. Discrete Applied Mathematics, 157(10):2425–2431.

[9] Barkan, C. L., Kelley, D. B., and Zornik, E. (2018). Premotor neuron divergence reflects vocal evolution. J Neurosci, 38(23):5325–5337.

[10] Barker, A. J. (2021). Brains and speciation: Control of behavior. Curr Opin Neurobiol, 71:158–163.

[11] Barsotti, E., Correia, A., and Cardona, A. (2021). Neural architectures in the light of comparative connectomics. Curr Opin Neurobiol, 71:139–149.

[12] Bassett, D. S. and Bullmore, E. (2006). Small-world brain networks. The neuroscientist, 12(6):512–523.

[13] Bassett, D. S. and Bullmore, E. T. (2009). Human brain networks in health and disease. Current opinion in neurology, 22(4):340.

[14] Bassett, D. S., Greenfield, D. L., Meyer-Lindenberg, A., Weinberger, D. R., Moore, S. W., and Bullmore, E. T. (2010). Efficient physical embedding of topologically complex information processing networks in brains and computer circuits. PLoS Comput Biol, 6(4):e1000748.

[15] Bassett, D. S., Porter, M. A., Wymbs, N. F., Grafton, S. T., Carlson, J. M., and Mucha, P. J. (2013). Robust detection of dynamic community structure in networks. Chaos: An Interdisciplinary Journal of Nonlinear Science, 23(1):013142.

[16] Bassett, D. S. and Sporns, O. (2017). Network neuroscience. Nature neuroscience, 20(3):353–364.

[17] Bendesky, A. and Bargmann, C. I. (2011). Genetic contributions to behavioural diversity at the gene-environment interface. Nature Reviews Genetics, 12(12):809–820.

[18] Beul, S. F., Grant, S., and Hilgetag, C. C. (2015). A predictive model of the cat cortical connectome based on cytoarchitecture and distance. Brain Struct Funct, 220(6):3167–3184.

[19] Blondel, V. D., Guillaume, J.-L., Lambiotte, R., and Lefebvre, E. (2008). Fast unfolding of communities in large networks. Journal of statistical mechanics: theory and experiment, 2008(10):P10008.

[20] Bota, M., Sporns, O., and Swanson, L. W. (2015). Architecture of the cerebral cortical association connectome underlying cognition. Proc Natl Acad Sci USA, 112(16):E2093–E2101.

[21] Brandes, U. (2001). A faster algorithm for betweenness centrality. Journal of mathematical sociology, 25(2):163–177.

[22] Buckner, R. L. and Krienen, F. M. (2013). The evolution of distributed association networks in the human brain. Trends Cogn Sci, 17(12):648–665.

[23] Bullmore, E. and Sporns, O. (2012). The economy of brain network organization. Nat Rev Neurosci, 13(5):336–349.

[24] Burke, J. D. (1968). A brief history of the taxonomy of mammals. MCV/Q, Medical College of Virginia Quarterly, 4(2):77–80.

[25] Chiang, A.-S., Lin, C.-Y., Chuang, C.-C., Chang, H.-M., Hsieh, C.-H., Yeh, C.-W., Shih, C.-T., Wu, J.-J., Wang, G.-T., Chen, Y.-C., et al. (2011). Three-dimensional reconstruction of brain-wide wiring networks in drosophila at single-cell resolution. Curr Biol, 21(1):1–11.

[26] Chung, F. (1996). Spectral graph theory. fresno. Proceedings of the American Mathematical Society.

[27] Colizza, V., Flammini, A., Serrano, M. A., and Vespignani, A. (2006). Detecting rich-club ordering in complex networks. Nature physics, 2(2):110–115.

[28] Consortium, Z. et al. (2020). A comparative genomics multitool for scientific discovery and conservation. Nature, 587(7833):240.

[29] Darwin, C. (1959). On the origin of species. Routledge.

[30] Das, K. C. (2004). The laplacian spectrum of a graph. Computers & Mathematics with Applications, 48(5-6):715–724.

[31] de Lange, S., de Reus, M., and Van Den Heuvel, M. (2014). The laplacian spectrum of neural networks. Front Comput Neurosci, 7:189.

[32] de Lange, S. C., van den Heuvel, M. P., and de Reus, M. A. (2016). The role of symmetry in neural networks and their laplacian spectra. NeuroImage, 141:357–365.

[33] de Reus, M. A. and van den Heuvel, M. P. (2013). Rich club organization and intermodule communication in the cat connectome. J Neurosci, 33(32):12929–12939.

[34] Ding, Y., Berrocal, A., Morita, T., Longden, K. D., and Stern, D. L. (2016). Natural courtship song variation caused by an intronic retroelement in an ion channel gene. Nature, 536(7616):329–332.

[35] Ding, Y., Lillvis, J. L., Cande, J., Berman, G. J., Arthur, B. J., Long, X., Xu, M., Dickson, B. J., and Stern, D. L. (2019). Neural evolution of context-dependent fly song. Current biology, 29(7):1089–1099.

[36] Eigenbrod, O., Debus, K. Y., Reznick, J., Bennett, N. C., Sánchez-Carranza, O., Omerbašić, D., Hart, D. W., Barker, A. J., Zhong, W., Lutermann, H., et al. (2019). Rapid molecular evolution of pain insensitivity in multiple african rodents. Science, 364(6443):852–859.

[37] Faskowitz, J., Betzel, R. F., and Sporns, O. (2021). Edges in brain networks: Contributions to models of structure and function. arXiv preprint arXiv:2105.07069.

[38] Fortunato, S. and Barthelemy, M. (2007). Resolution limit in community detection. Proceedings of the national academy of sciences, 104(1):36–41.

[39] Freeman, L. C. (1978). Centrality in social networks conceptual clarification. Social networks, 1(3):215–239.

[40] Grone, R. and Merris, R. (1994). The laplacian spectrum of a graph ii. SIAM Journal on discrete mathematics, 7(2):221–229.

[41] Grone, R., Merris, R., and Sunder, V. (1990). The laplacian spectrum of a graph. SIAM Journal on matrix analysis and applications, 11(2):218–238.

[42] Hernández-Hernández, T., Miller, E. C., Román-Palacios, C., and Wiens, J. J. (2021). Speciation across the t ree of l ife. Biological Reviews, 96(4):1205–1242.

[43] Hilgetag, C. C. and Kaiser, M. (2004). Clustered organization of cortical connectivity. Neuroinformatics, 2(3):353–360.

[44] Humphries, M. D. and Gurney, K. (2008). Network ‘small-world-ness’: a quantitative method for determining canonical network equivalence. PloS one, 3(4):e0002051.

[45] Insel, T. R. and Shapiro, L. E. (1992). Oxytocin receptor distribution reflects social organization in monogamous and polygamous voles. Proceedings of the National Academy of Sciences, 89(13):5981–5985.

[46] Jaggard, J. B., Lloyd, E., Yuiska, A., Patch, A., Fily, Y., Kowalko, J. E., Appelbaum, L., Duboue, E. R., and Keene, A. C. (2020). Cavefish brain atlases reveal functional and anatomical convergence across independently evolved populations. Sci Adv, 6(38):eaba3126.

[47] Khallaf, M. A., Auer, T. O., Grabe, V., Depetris-Chauvin, A., Ammagarahalli, B., Zhang, D.-D., Lavista-Llanos, S., Kaftan, F., Weißflog, J., Matzkin, L. M., et al. (2020). Mate discrimination among subspecies through a conserved olfactory pathway. Sci Adv, 6(25):eaba5279.

[48] Kintali, S. (2008). Betweenness centrality: Algorithms and lower bounds. arXiv preprint arXiv:0809.1906.

[49] Latora, V. and Marchiori, M. (2001). Efficient behavior of small-world networks. Physical review letters, 87(19):198701.

[50] Leemans, A., Jeurissen, B., Sijbers, J., and Jones, D. (2009). Exploredti: a graphical toolbox for processing, analyzing, and visualizing diffusion mr data. In Proc Intl Soc Mag Reson Med, volume 17, page 3537.

[51] Leung, C. and Chau, H. (2007). Weighted assortative and disassortative networks model. Physica A: Statistical Mechanics and its Applications, 378(2):591–602.

[52] Liu, Z.-Q., Zheng, Y.-Q., and Misic, B. (2020). Network topology of the marmoset connectome. Net Neurosci, 4(4):1181–1196.

[53] Loomis, C., Peuß, R., Jaggard, J. B., Wang, Y., McKinney, S. A., Raftopoulos, S. C., Raftopoulos, A., Whu, D., Green, M., McGaugh, S. E., et al. (2019). An adult brain atlas reveals broad neuroanatomical changes in independently evolved populations of mexican cavefish. Frontiers in neuroanatomy, 13:88.

[54] Maier-Hein, K. H., Neher, P. F., Houde, J.-C., Côté, M.-A., Garyfallidis, E., Zhong, J., Chamberland, M., Yeh, F.-C., Lin, Y.-C., Ji, Q., et al. (2017). The challenge of mapping the human connectome based on diffusion tractography. Nat Commun, 8(1):1–13.

[55] Majka, P., Chaplin, T. A., Yu, H.-H., Tolpygo, A., Mitra, P. P., Wójcik, D. K., and Rosa, M. G. (2016). Towards a comprehensive atlas of cortical connections in a primate brain: Mapping tracer injection studies of the common marmoset into a reference digital template. J Com Neurol, 524(11):2161–2181.

[56] Mallet, J. (1995). A species definition for the modern synthesis. Trends in Ecology & Evolution, 10(7):294–299.

[57] Markov, N. T., Ercsey-Ravasz, M., Ribeiro Gomes, A., Lamy, C., Magrou, L., Vezoli, J., Misery, P., Falchier, A., Quilodran, R., Gariel, M., et al. (2012). A weighted and directed interareal connectivity matrix for macaque cerebral cortex. Cereb Cortex, 24(1):17–36.

[58] Markow, T. A. and O’Grady, P. M. (2005). Evolutionary genetics of reproductive behavior in drosophila: connecting the dots. Annu. Rev. Genet., 39:263–291.

[59] Mars, R. B., Jbabdi, S., and Rushworth, M. F. (2021). A common space approach to comparative neuroscience. Annual Review of Neuroscience, 44.

[60] Mars, R. B., Passingham, R. E., and Jbabdi, S. (2018a). Connectivity fingerprints: from areal descriptions to abstract spaces. Trends in cognitive sciences, 22(11):1026–1037.

[61] Mars, R. B., Sotiropoulos, S. N., Passingham, R. E., Sallet, J., Verhagen, L., Khrapitchev, A. A., Sibson, N., and Jbabdi, S. (2018b). Whole brain comparative anatomy using connectivity blueprints. Elife, 7:e35237.

[62] Mars, R. B., Verhagen, L., Gladwin, T. E., Neubert, F.-X., Sallet, J., and Rushworth, M. F. (2016). Comparing brains by matching connectivity profiles. Neuroscience & Biobehavioral Reviews, 60:90–97.

[63] Martinez, P. and Sprecher, S. G. (2020). Of circuits and brains: The origin and diversification of neural architectures. Frontiers in Ecology and Evolution, 8:82.

[64] Maslov, S. and Sneppen, K. (2002). Specificity and stability in topology of protein networks. Science, 296(5569):910–913.

[65] McAuley, J. J., da Fontoura Costa, L., and Caetano, T. S. (2007). Rich-club phenomenon across complex network hierarchies. Applied Physics Letters, 91(8):084103.

[66] Mišić, B. and Sporns, O. (2016). From regions to connections and networks: new bridges between brain and behavior. Curr Opin Neurobiol, 40:1–7.

[67] Murphy, W. J., Foley, N. M., Bredemeyer, K. R., Gatesy, J., and Springer, M. S. (2021). Phylogenomics and the genetic architecture of the placental mammal radiation. Annual Review of Animal Biosciences, 9:29–53.

[68] Newman, M. E. (2002). Assortative mixing in networks. Physical review letters, 89(20):208701.

[69] Newman, M. E. (2003). The structure and function of complex networks. SIAM review, 45(2):167–256.

[70] Newman, M. W. (2001). The laplacian spectrum of graphs.

[71] Oh, S. W., Harris, J. A., Ng, L., Winslow, B., Cain, N., Mihalas, S., Wang, Q., Lau, C., Kuan, L., Henry, A. M., et al. (2014). A mesoscale connectome of the mouse brain. Nature, 508(7495):207–214.

[72] Onnela, J.-P., Saramäki, J., Kertész, J., and Kaski, K. (2005). Intensity and coherence of motifs in weighted complex networks. Physical Review E, 71(6):065103.

[73] O’Grady, P. M. and DeSalle, R. (2018). Phylogeny of the genus drosophila. Genetics, 209(1):1–25.

[74] Pantoja, C., Larsch, J., Laurell, E., Marquart, G., Kunst, M., and Baier, H. (2020). Rapid effects of selection on brain-wide activity and behavior. Current Biology, 30(18):3647–3656.

[75] Park, T. J., Lu, Y., Jüttner, R., Smith, E. S. J., Hu, J., Brand, A., Wetzel, C., Milenkovic, N., Erdmann, B., Heppenstall, P. A., et al. (2008). Selective inflammatory pain insensitivity in the african naked mole-rat (heterocephalus glaber). PLoS biology, 6(1):e13.

[76] Passingham, R. E., Stephan, K. E., and Kötter, R. (2002). The anatomical basis of functional localization in the cortex. Nat Rev Neurosci, 3(8):606–616.

[77] Rubinov, M. and Sporns, O. (2010). Complex network measures of brain connectivity: uses and interpretations. Neuroimage, 52(3):1059–1069.

[78] Rubinov, M., Ypma, R. J., Watson, C., and Bullmore, E. T. (2015). Wiring cost and topological participation of the mouse brain connectome. Proc Natl Acad Sci USA, 112(32):10032–10037.

[79] Scannell, J. W., Blakemore, C., and Young, M. P. (1995). Analysis of connectivity in the cat cerebral cortex. J Neurosci, 15(2):1463–1483.

[80] Schilling, K. G., Nath, V., Hansen, C., Parvathaneni, P., Blaber, J., Gao, Y., Neher, P., Aydogan, D. B., Shi, Y., Ocampo-Pineda, M., et al. (2019). Limits to anatomical accuracy of diffusion tractography using modern approaches. NeuroImage, 185:1–11.

[81] Seehausen, O., Butlin, R. K., Keller, I., Wagner, C. E., Boughman, J. W., Hohenlohe, P. A., Peichel, C. L., Saetre, G.-P., Bank, C., Brännstrüm, A., et al. (2014). Genomics and the origin of species. Nat Rev Genet, 15(3):176–192.

[82] Seeholzer, L. F., Seppo, M., Stern, D. L., and Ruta, V. (2018). Evolution of a central neural circuit underlies drosophila mate preferences. Nature, 559(7715):564–569.

[83] Shanahan, M., Bingman, V. P., Shimizu, T., Wild, M., and Güntürkün, O. (2013). Large-scale network organization in the avian forebrain: a connectivity matrix and theoretical analysis. Front Comput Neurosci, 7:89.

[84] Smith, E. S. J., Omerbašić, D., Lechner, S. G., Anirudhan, G., Lapatsina, L., and Lewin, G. R. (2011). The molecular basis of acid insensitivity in the african naked mole-rat. Science, 334(6062):1557–1560.

[85] Sporns, O. (2013). Network attributes for segregation and integration in the human brain. Curr Opin Neurobiol, 23(2):162–171.

[86] Sporns, O., Tononi, G., and Kötter, R. (2005). The human connectome: a structural description of the human brain. PLoS Comput Biol, 1(4):e42.

[87] Sporns, O. and Zwi, J. D. (2004). The small world of the cerebral cortex. Neuroinformatics, 2(2):145–162.

[88] Stiso, J. and Bassett, D. S. (2018). Spatial embedding imposes constraints on neuronal network architectures. Trends Cogn Sci, 22(12):1127–1142.

[89] Suárez, L. E., Markello, R. D., Betzel, R. F., and Misic, B. (2020). Linking structure and function in macroscale brain networks. Trends Cogn Sci, 24(4):302–315.

[90] Suárez, L. E., Richards, B. A., Lajoie, G., and Misic, B. (2021). Learning function from structure in neuromorphic networks. Nature Machine Intelligence, 3(9):771–786.

[91] Tendler, B. C., Hanayik, T., Ansorge, O., Bangerter-Christensen, S., Berns, G. S., Bertelsen, M. F., Bryant, K. L., Foxley, S., Howard, A. F., Huszar, I., et al. (2021). The digital brain bank: an open access platform for postmortem datasets. bioRxiv.

[92] Tononi, G., Sporns, O., and Edelman, G. M. (1994). A measure for brain complexity: relating functional segregation and integration in the nervous system. Proceedings of the National Academy of Sciences, 91(11):5033–5037.

[93] Tournier, J.-D., Calamante, F., Gadian, D. G., and Connelly, A. (2004). Direct estimation of the fiber orientation density function from diffusion-weighted mri data using spherical deconvolution. NeuroImage, 23(3):1176–1185.

[94] Towlson, E. K., Vértes, P. E., Ahnert, S. E., Schafer, W. R., and Bullmore, E. T. (2013). The rich club of the c. elegans neuronal connectome. J Neurosci, 33(15):6380–6387.

[95] Van den Heuvel, M. P., Bullmore, E. T., and Sporns, O. (2016). Comparative connectomics. Trends Cogn Sci, 20(5):345–361.

[96] Van Den Heuvel, M. P., Kahn, R. S., Goñi, J., and Sporns, O. (2012). High-cost, high-capacity backbone for global brain communication. Proc Natl Acad Sci, 109(28):11372–11377.

[97] van den Heuvel, M. P., Mandl, R. C., Stam, C. J., Kahn, R. S., and Pol, H. E. H. (2010). Aberrant frontal and temporal complex network structure in schizophrenia: a graph theoretical analysis. Journal of Neuroscience, 30(47):15915–15926.

[98] Van Den Heuvel, M. P. and Sporns, O. (2011). Rich-club organization of the human connectome. Journal of Neuroscience, 31(44):15775–15786.

[99] Vanwalleghem, G. C., Ahrens, M. B., and Scott, E. K. (2018). Integrative whole-brain neuroscience in larval zebrafish. Current opinion in neurobiology, 50:136–145.

[100] Warrington, S., Thompson, E., Bastiani, M., Dubois, J., Baxter, L., Slater, R., Jbabdi, S., Mars, R., and Sotiropoulos, S. (2022). Concurrent mapping of brain ontogeny and phylogeny within a common connectivity space. bioRxiv.

[101] Watts, D. J. and Strogatz, S. H. (1998). Collective dynamics of ‘small-world’ networks. Nature, 393(6684):440–442.

[102] White, J. G., Southgate, E., Thomson, J. N., Brenner, S., et al. (1986). The structure of the nervous system of the nematode caenorhabditis elegans. Philos Trans R Soc Lond B Biol Sci, 314(1165):1–340.

[103] Winslow, J. T., Hastings, N., Carter, C. S., Harbaugh, C. R., and Insel, T. R. (1993). A role for central vasopressin in pair bonding in monogamous prairie voles. Nature, 365(6446):545–548.

[104] Worrell, J. C., Rumschlag, J., Betzel, R. F., Sporns, O., and Mišić, B. (2017). Optimized connectome architecture for sensory-motor integration. Net Neurosci, 1(4):415–430.

[105] Yokoyama, C., Autio, J. A., Ikeda, T., Sallet, J., Mars, R. B., Van Essen, D. C., Glasser, M. F., Sadato, N., and Hayashi, T. (2021). Comparative connectomics of the primate social brain. NeuroImage, 245:118693.

[106] York, R. A. (2018). Assessing the genetic landscape of animal behavior. Genetics, 209(1):223–232.

[107] York, R. A., Byrne, A., Abdilleh, K., Patil, C., Streelman, T., Finger, T. E., and Fernald, R. D. (2019). Behavioral evolution contributes to hindbrain diversification among lake malawi cichlid fish. Scientific reports, 9(1):1–9.

[108] Zamora-López, G., Zhou, C., and Kurths, J. (2010). Cortical hubs form a module for multisensory integration on top of the hierarchy of cortical networks. Front Neuroin-form, 4:1.

[109] Zhou, S. and Mondragón, R. J. (2004). The rich-club phenomenon in the internet topology. IEEE communications letters, 8(3):180–182.

